# ROCK2 inhibition has a dual role in reducing ECM remodelling and cell growth, while impairing migration and invasion

**DOI:** 10.1101/2025.07.21.666015

**Authors:** Daniel A. Reed, Anna E. Howell, Nadia Kuepper, Alice M. H. Tran, Astrid Magenau, Deborah S. Barkauskas, Max Nobis, Cecilia R. Chambers, Victoria Lee, Lily M. Channon, Jessie Zhu, Shona Ritchie, Janett Stoehr, Kaitlin Wylie, Julia Chen, Denise Attwater, Kate Harvey, Sunny Z. Wu, Kate Saw, Ruth J. Lyons, Anaiis Zaratzian, Michael Tayao, Andrew Da Silva, David Gallego-Ortega, Anthony J. Gill, Thomas R. Cox, Brooke A. Pereira, Kendelle J. Murphy, Jennifer P. Morton, Elgene Lim, Alexander Swarbrick, Sandra O’Toole, Michael S. Samuel, C. Elizabeth Caldon, Alexandra Zanin-Zhorov, Paul Timpson, David Herrmann

## Abstract

The Rho-associated coiled-coil containing protein kinases 1/2 (ROCK1/2) are key signalling proteins involved in the regulation of the actin cytoskeleton and control a variety of cellular processes. This includes cell proliferation, stemness, cell migration and invasion as well as actomyosin contraction and extracellular matrix (ECM) remodelling. Due to a lack of ROCK2 specific inhibitors, previous pharmacological studies of ROCK function have relied on pan-ROCK1/2 inhibition. Here, we use the ROCK2 specific inhibitor, GV101, in the context of breast cancer (BC) models to uncouple the effects of ROCK2 pharmacological inhibition on both epithelial and stromal fibroblast cell populations. We demonstrate that ROCK2 inhibition with GV101 reduces fibroblast-mediated remodelling of pre-existing and *de novo* synthesised ECM resulting in a reduction in biomechanical stiffness, while also limiting epithelial cell growth in 2D and 3D settings. We also demonstrate that ROCK2 inhibition exposes epithelial cell vulnerability to fluid flow-induced shear stress. Furthermore, using 3D co-cultures we reveal that GV101 reduces single-cell migratory capacity, while the disruption of stromal ECM architecture is sufficient to impede collective cell migration and invasion. Finally, we assessed ROCK2 expression in human BC patient tumours revealing that ROCK2 expression is up-regulated during BC progression and that high ROCK2 expression correlates with poor patient outcome in the triple-negative subtype of BC. Together, these results demonstrate that ROCK2 specific inhibition disrupts key functions in both epithelial cell and stromal fibroblast compartments warranting further assessment of ROCK2 in highly proliferative, fibrotic or metastatic diseases, such as BC.

**Highlights:**

- Fibroblast-mediated remodelling of pre-existing and *de novo* synthesised extracellular matrix is reduced upon ROCK2 specific targeting.
- ROCK2 inhibition impairs epithelial cancer cell growth and colony formation in 2D and 3D contexts, while also limiting single-cell migration and cell survival upon fluid flow-induced shear stress.
- ROCK2 targeting during fibroblast-mediated ECM remodelling is sufficient to disrupt collective cell migration as well as 3D invasion into organotypic fibroblast-contracted collagen matrices.
- ROCK2 expression is up-regulated during breast cancer progression and high expression correlates with poor patient outcome in triple-negative breast cancer.

## Introduction

The extracellular environment delivers both mechanical and biochemical signals that influence cellular interactions with their surroundings, leading to alterations in various cellular functions during development, tissue homeostasis and disease progression. The Rho-associated coiled-coil containing protein kinases 1/2 (ROCK1/2) are central regulators of actin cytoskeletal processes, integrating multiple cellular processes that are crucial for both developmental and homeostatic physiology. Activation of ROCK predominantly occurs through binding of the small GTPase RhoA, a key signalling node facilitating outside-in-signalling from multiple extracellular inputs, resulting in a conformational change and subsequent relieving of auto-inhibitory interaction [1]. ROCK activation results in the phosphorylation of multiple downstream targets including myosin II regulatory light chain kinase (MLCK), the myosin binding subunit MYPT1 and LIM kinases, which can further regulate actin-binding proteins, such as cofilin [2]. This leads to the collective promotion of actin filament stabilisation and bundling, driving actomyosin contraction [3–5]. Due to its crucial role in actin cytoskeleton regulation, ROCK signalling is critical for a variety of cytoskeleton-dependent cellular processes including cell proliferation [6–8], survival and apoptosis [9–11], migration and invasion [12–15], anchorage-independent growth [16–18], anoikis [19] and resistance to shear stress forces [20, 21]. Moreover, extracellular matrix (ECM) deposition and remodelling leading to the generation of a biomechanically stiff environment can be promoted further by ROCK activation [22, 23]. As a result, the ROCK signalling axis has been implicated in a variety of diseases and hence inhibition of ROCK has long been pursued as a possible therapeutic option in multiple pathologies [1, 24, 25].

ROCK1/2 are widely expressed in embryonic and adult tissues [26] and while the two proteins have many overlapping functions, they often exhibit tissue, cell or even subcellular specific roles [27]. Genetic studies have been critical in understanding the individual roles of both ROCK1 and ROCK2 in homeostatic and disease contexts, while pharmacological studies have been limited to the use of pan-ROCK1/2 inhibitors such as Fasudil and Y27632. We and others have shown that these ROCK1/2 inhibitors demonstrate promising proof-of-principle efficacy of ROCK inhibition in a variety of pre-clinical disease contexts, including cardiovascular disease [28], neurodegenerative disorders [29, 30], autoimmune disease [31, 32], idiopathic pulmonary fibrosis [33, 34] and various solid cancers [35–39]. However, the translation of these findings has been difficult due to the low specificity and tolerability of pan-ROCK1/2 inhibitors [40].

ROCK2 specific inhibitors therefore offer the promise of on-target ROCK2 inhibition while avoiding the negative side effects associated with previous pan ROCK1/2 inhibitors. For example, Belumosudil has shown promising results for the treatment of fibrotic diseases [41–43] and immunological disorders [44–47], yet still exhibits off-target kinase inhibition [46, 48], limiting its evaluation on targeting the ROCK2 signalling pathway specifically. Excitingly, a more potent ROCK2 inhibitor, GV101 (1000-fold selective for the ROCK2 isoform over ROCK1) was recently shown to stall fibrosis in a mouse model of non-alcoholic fatty liver disease [49]. However, the effects of ROCK2 inhibitors on basic cellular function, such as cell proliferation, migration, invasion and ECM remodelling, have not yet been assessed in other disease settings. This therefore warrants further investigation to uncover the effects of ROCK2 inhibition on these key cellular processes.

Here we use triple-negative breast cancer (TNBC), a disease characterised by highly proliferative and invasive epithelial cells as well as significant ECM deposition and remodelling [50, 51], as a model system to decipher the effects of ROCK2 inhibition using GV101 on epithelial and stromal cell populations in both murine and human-derived settings. We demonstrate that ROCK2 inhibition reduces fibroblast-mediated remodelling of pre-existing and *de novo* synthesised collagen as well as decreasing epithelial cell growth and colony formation in 2-dimensional (2D) contexts and under 3D anchorage-independent growth settings. We further investigated the effects of ROCK2 inhibition on epithelial cell migration on cell-derived matrices (CDMs) and show that GV101 reduces the single-cell migratory capacity of epithelial cells, while disrupting the matrix production by fibroblast cells with GV101 altered subsequent epithelial collective cell migration. We also reveal that ROCK2 inhibition following fluid flow-induced shear stress enhances vulnerability to cell death. Moreover, we uncouple the effects of epithelial *versus* stromal ROCK2 targeting by assessing epithelial cell invasion into 3D fibroblast-contracted organotypic matrices, where ROCK2 inhibition to disrupt collagen remodelling is sufficient to inhibit epithelial cell invasion. Finally, we provide evidence that ROCK2 may play an important role in disease progression and survival in the triple-negative subtype of breast cancer, suggesting that ROCK2 inhibition should be further investigated in this highly lethal disease.

## Results

### ROCK2 inhibition with GV101 reduces fibroblast-mediated remodelling of pre-existing and *de novo* ECM

The role of the ROCK signalling axis in ECM remodelling and stiffening has been well documented [6, 22, 35, 36]. Conversely, RhoA-ROCK signalling can also be activated by ECM tension and rigid environments [22, 37]. This leads to further ECM and tissue stiffening, driving a feed-forward loop that contributes to the pathogenesis of various diseases involving tissue fibrosis [52–57]. Here, we first investigated how the ROCK2 specific inhibitor GV101 could modulate ECM remodelling in the setting of TNBC, a highly fibrotic BC subtype [50]. One of the key cell types involved in ECM synthesis and remodelling in tumours are cancer-associated fibroblasts (CAFs), which are present in mouse models of the disease and human patients [58, 59]. In this study we utilised CAFs previously isolated from end-stage tumours of the genetically engineered MMTV-PyMT mouse model of BC [37, 60] as well as CAFs isolated from human TNBC patients [58, 61]. Both PyMT and human CAFs were shown to express ROCK2 (Figure S1A) and phosphorylation of its downstream target Cofilin [2, 49], which was decreased upon GV101 treatment (Figure S1B and C). This data indicates that we can target ROCK2 in the stromal TNBC fibroblast compartment as previously achieved in the context of liver fibrosis [49].

To investigate the effects of ROCK2 inhibition *via* GV101 on CAF-mediated collagen contraction and remodelling, we initially performed organotypic contraction assays, which provide a malleable platform from which to assess key features of CAF-mediated contraction, remodelling and biomechanics of pre-existing ECM [62]. Here, CAFs were embedded into acid-extracted fibrillar collagen and allowed to contract and remodel the collagen matrix over a 12-day period, as previously described (Figure 1A) [35, 63–66]. Treatment with either GV101 or Fasudil during collagen contraction reduced the contractile properties of PyMT CAFs compared to vehicle control, as seen by a significant increase in matrix size at day 12 (Figure 1B). Fasudil treatment resulted in an additional decrease in contraction compared to GV101 (Figure 1B), showing that both ROCK2 specific and pan ROCK1/2 inhibitors can reduce ECM contraction. In order to assess the biomechanical properties of the organotypic matrix, we performed unconfined compression analysis, as previously achieved [65, 66]. This showed that both GV101 and Fasudil treated matrices exhibited a significant reduction in organotypic matrix stiffness (Young’s modulus) compared to vehicle control (Figure 1D). Extracellular tension and stiff environments can further promote ROCK signalling through RhoA activation, and hence GV101 may be able to inhibit this feed-forward loop by reducing both ROCK2 signalling as well as its concomitant activation from the environment.

**Figure 1:**
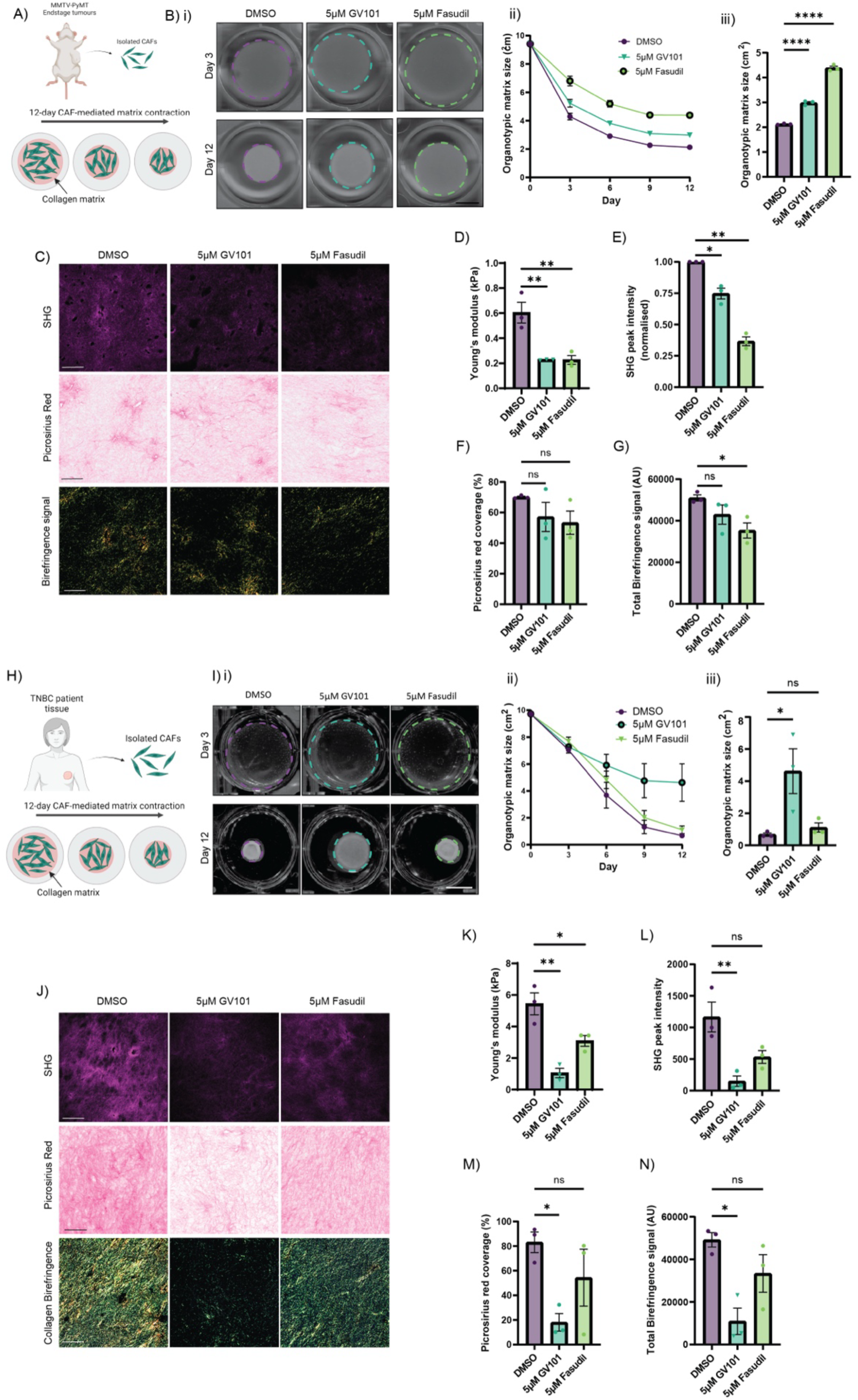
ROCK2 inhibition with GV101 reduces CAF-mediated remodelling of pre-existing collagen in murine and human models of TNBC, also see Figure S1. **(A)** Schematic of collagen contraction assays utilising CAFs isolated from end-stage MMTV-PyMT tumours. **(B)** Representative images of PyMT CAF matrices treated with DMSO, GV101 or Fasudil at days 3 and 12 of contraction (i) and quantification of PyMT CAF organotypic matrix size over a 12-day contraction period (ii) and on day 12 (iii). **(C)** Representative SHG images of PyMT CAF matrices to assess collagen I (top panel, magenta), picrosirius red staining to assess fibrillar collagen (middle panel) and polarised light birefringence imaging to assess collagen bundling and density (bottom panel). **(D)** Quantification of unconfined compression analysis to assess Young’s modulus. **(E-G)** Quantification of PyMT CAF matrices; SHG peak signal intensity **(E)**, picrosirius red staining coverage **(F)** and total birefringence signal **(G). (H)** Schematic of collagen contraction assays utilising CAFs isolated from human TNBC patients. **(I)** Representative images of human CAF matrices treated with DMSO, GV101 or Fasudil at days 3 and 12 of contraction **(i)** and quantification of CAF organotypic matrix size over a 12-day contraction period (ii) and on day 12 (iii). (J) Representative SHG images of human CAF matrices to assess collagen I (top panel, magenta), picrosirius red staining to assess fibrillar collagen (middle panel) and polarised light birefringence imaging to assess collagen bundling and density (bottom panel). **(K)** Quantification of unconfined compression analysis to assess matrix Young’s modulus. **(L-N)** Quantification of human CAF matrices; SHG peak signal intensity **(L)**, picrosirius red staining coverage (M) and total birefringence signal **(N)**. Data represents mean ± SEM of three individual repeats. p-values were determined using One-sample t and Wilcoxon test for normalised data and for remaining data a One-way ANOVA with multiple comparisons was used. ns = not significant (p≥0.05), * = p<0.05, ** = p<0.01, *** = p<0.001, **** = p<0.0001. Scale bar = 1cm (B and I), 100µm (C and J).

We next employed multiphoton-based second harmonic generation (SHG) imaging and picrosirius red staining to evaluate how GV101 affected the abundance and remodelling of fibrillar collagen. SHG imaging of fibrillar collagen I content (Figure 1C, top panel, magenta and Movie S1) demonstrated a significant decrease in SHG signal intensity in both GV101 and Fasudil treated matrices compared to vehicle control (Figure 1E). Moreover, picrosirius red staining to assess total fibrillar collagen I/III content (Figure 1C, middle panel) revealed a subtle, non-significant decrease in picrosirius red staining coverage (Figure 1F). We also performed polarised light imaging of the picrosirius red stained matrices to assess collagen birefringence where we observed a subtle decrease in total birefringence signal (Figure 1G). Interestingly, analysis of green, yellow and red-orange birefringence signal (indicative of low, intermediate and highly bundled collagen fibres respectively) showed a significant decrease in red-orange birefringence signal (Figure S1D and G) upon GV101 treatment. This correlates to a decrease in highly bundled collagen fibres following ROCK2 inhibition and suggesting that ROCK2 may be required for the generation of mature collagen fibres in this setting. Treatment with Fasudil reduced green, yellow and red-orange birefringence signal (Figure S1D to G, respectively), as well as the total birefringence signal (Figure 1G), suggesting that pan ROCK1/2 inhibitors can limit early and late remodelling of collagen/ECM. Collectively, these data show that GV101-mediated ROCK2 inhibition impairs CAF function on pre-existing ECM at several levels including CAF-mediated collagen matrix remodelling, contraction and stiffening.

In order to corroborate our findings from mouse PyMT CAFs, we next assessed the effects of GV101 on human TNBC CAF function. As before, human TNBC CAFs [58, 61] were embedded into fibrillar collagen for matrix contraction (Figure 1H). Similar to the murine model, human TNBC CAFs showed a significantly decreased ability to contract the collagen matrix when treated with GV101 compared to vehicle (Figure 1I). Interestingly, in this setting no significant differences in contraction were observed when CAF matrices were treated with Fasudil compared to vehicle (Figure 1I). Assessment of the biomechanical properties of these matrices similarly showed a significant decrease in organotypic matrix stiffness upon GV101-mediated ROCK2 inhibition (Figure 1K). Multiphoton imaging showed a decrease in SHG signal intensity indicative of lower fibrillar collagen I content in GV101-treated matrices (Figure 1J, top panel, Figure 1L and Movie S2), while picrosirius red staining (Figure 1J, middle panel) revealed a decrease in total fibrillar collagen I/III coverage in the matrices with GV101 treatment (Figure 1M). Moreover, polarised light birefringence imaging demonstrated a reduction in total (Figure 1N), green (Figure S1H and I), yellow (Figure S1H and J) and red-orange (Figure S1H and K) birefringence signal upon GV101 treatment, suggesting that ROCK2 inhibition in the human TNBC CAF context affects all stages of collagen fibre maturation, bundling and crosslinking. Conversely, Fasudil treatment only resulted in a significant decrease in matrix stiffness (Figure 1K) with a trend towards decreased SHG (Figure 1L), picrosirius red coverage (Figure 1M) and birefringence signal (Figure 1N, Figure S1H to K). Overall, these data using human TNBC CAFs align with the data obtained in the murine PyMT CAF setting showing that GV101 can inhibit fibroblast-mediated collagen remodelling, contraction and stiffness.

While the function of ROCK2 to facilitate the remodelling of pre-existing ECM has been characterised, less is known about the effects of ROCK2 inhibition on the *de novo* production of ECM. In order to assess this aspect of CAF biology in relation to ROCK2 inhibition, cell-derived matrix (CDM) assays were employed as described previously [35, 65, 67–69]. Here CAFs are grown in a monolayer and then treated with ascorbic acid to stimulate matrix production with concurrent ROCK2 or pan-ROCK1/2 inhibition [35, 65, 68] (Figure 2A). SHG imaging was then used to characterise *de novo* deposited ECM (Figure 2B, top panel and Movie S3 and Movie S4). CDMs generated by PyMT CAFs treated with either GV101 or Fasudil, showed a significant decrease in SHG signal intensity compared to vehicle control indicative of a reduction in fibrillar collagen I abundance (Figure 2Ci). Furthermore, brightfield imaging of picrosirius red stained CDMs confirmed a decrease in fibrillar collagen production across both treatment groups compared to vehicle (Figure 2B, bottom panel and Figure 2Cii), corroborating that GV101-mediated ROCK2 inhibition impairs the deposition and remodelling of *de novo* synthesised fibrillar collagen. Similarly, in the human TNBC CAF setting we observe that GV101 reduced SHG signal intensity and picrosirius red coverage to the same extent as Fasudil, as observed before in murine CAFs (Figure 2D and E and Movie S5 and Movie S6). Collectively, this data demonstrates a requirement for ROCK2 in CAF biology, in particular the contraction and remodelling of pre-existing collagen as well as the deposition and remodelling of *de novo* synthesised ECM in both murine and human breast cancer CAFs.

**Figure 2:**
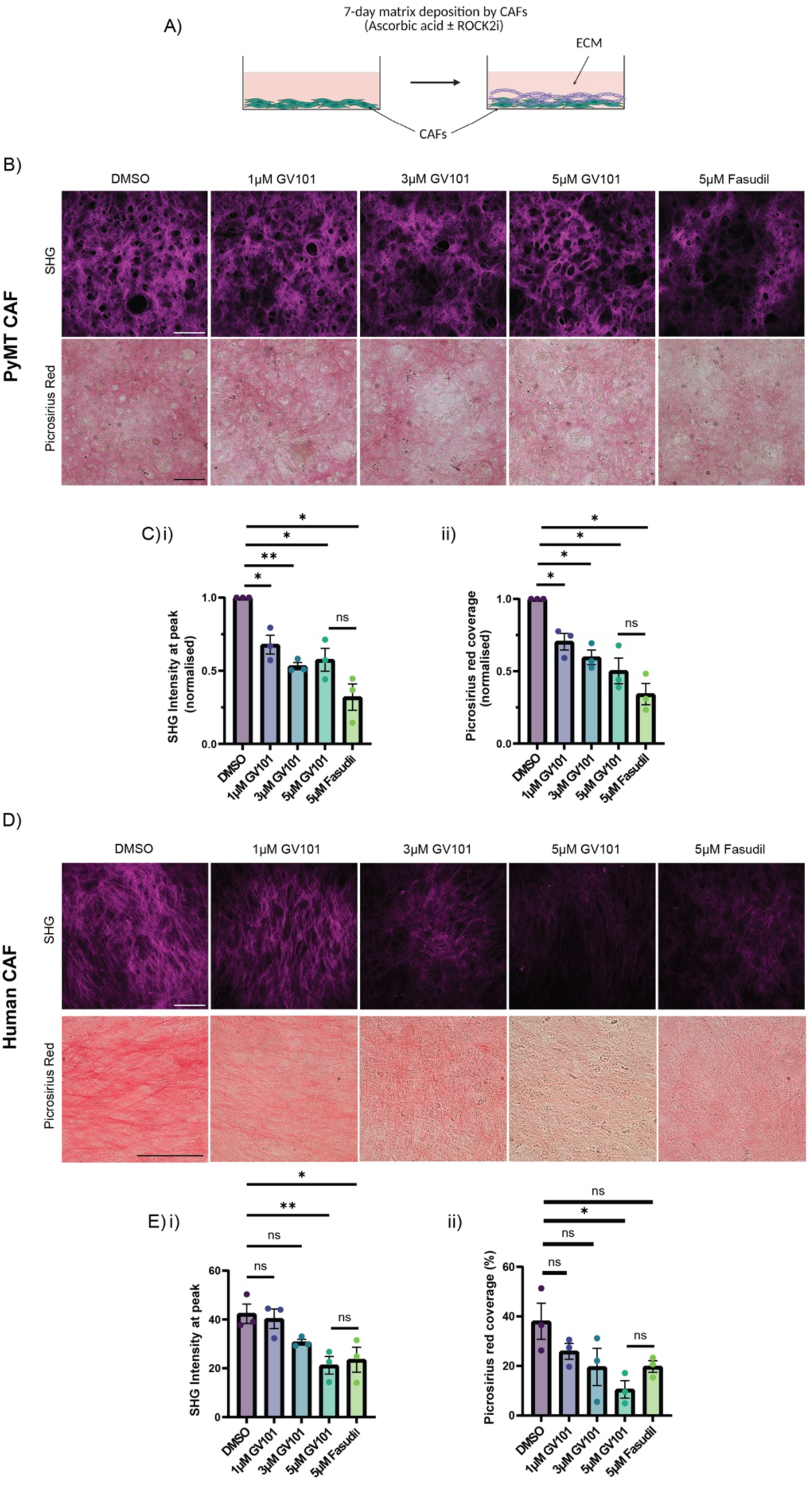
GV101 inhibits CAF-mediated remodelling of *de novo* synthesised collagen in cell-derived matrix assays. **(A)** Schematic of cell-derived matrix (CDM) production assays where both PyMT and human CAFs are treated with ascorbic acid to stimulate matrix production in the presence of DMSO, GV101 or Fasudil. **(B)** Representative images of PyMT CAF derived CDMs showing SHG imaging (top panel, magenta) and picrosirius red stained matrices (bottom panel). **(C)** Quantification of peak SHG signal (i) and picrosirius red staining coverage for PyMT CAF generated CDMs (ii). **(D)** Representative images of human CAF derived CDMs showing SHG imaging (top panel, magenta) and picrosirius red stained matrices (bottom panel). **(E)** Quantification of peak SHG signal (i) and picrosirius red coverage (ii) for human CAF generated CDMs. Data represents mean ± SEM of three individual repeats. p-values were determined using One-sample t and Wilcoxon test for normalised data and for remaining data a One-way ANOVA with multiple comparisons was used. ns = not significant (p≥0.05), * = p<0.05, ** = p<0.01. Scale bar = 100μm.

### ROCK2 inhibition with GV101 reduces epithelial cell growth in 2D and 3D and sensitises cells to fluid flow-induced shear stress

In addition to its role in stromal populations, ROCK signalling has also previously been identified to promote pathways involved in cell growth and survival [7, 9, 11]. Here, we assessed ROCK2 inhibition on these cellular functions utilising murine BC cells, isolated from end-stage tumours of the genetically engineered MMTV-PyMT mouse model [60, 70], and the human TNBC cell line MDA-MB-231. Both cell lines demonstrated ROCK2 expression (Figure S2A) and as shown before in CAFs (Figure S1), treatment of both cell lines with GV101 showed a decrease in pCofilin (Figure S2B and C), demonstrating that GV101 is able to reduce ROCK2 signalling in these cells. We first assessed the effects of GV101 on baseline cell viability. Here, both murine PyMT and human MDA-MB-231 cells treated with GV101 showed a significant decrease in cell viability compared to vehicle control with only a subtle decrease in cell viability observed using the pan-ROCK1/2 inhibitor Fasudil (Figure S2D and E) as previously shown in other contexts [35, 38, 39].

Next, we aimed to assess the effect of ROCK2 inhibition on colony outgrowth from individual cells using 2D colony formation assays (Figure 3A and B). Here, GV101 treatment significantly reduced the colony forming capacity of PyMT cancer cells at all concentrations tested compared to vehicle (Figure 3Ai and ii). Similar growth inhibition effects were observed in the human MDA-MB-231 TNBC cell line where GV101 treatment at all concentrations tested was sufficient to reduce colony outgrowth (Figure 3Bi and ii). We next investigated the effect of ROCK2 inhibition on 3D colony survival and outgrowth under AIG conditions [16, 18]. Here, single PyMT and MDA-MB-231 cancer cells were suspended in agarose and incubated for 14 and 10 days, respectively, as previously described [35, 65]. Both PyMT (Figure 3C) and MDA-MB-231 cells (Figure 3E) treated with GV101 showed a reduced ability to form 3D AIG clusters, as seen by a significant decrease in the number of clusters (Figure 3Di and Fi), as well as a decrease in the size of colonies that are able to grow (Figure 3Dii and Fii), indicative of both a reduction in 3D cell survival and proliferation, respectively. Collectively, these data demonstrate that ROCK2 inhibition can affect colony outgrowth from individual cells under both 2D and 3D conditions.

**Figure 3:**
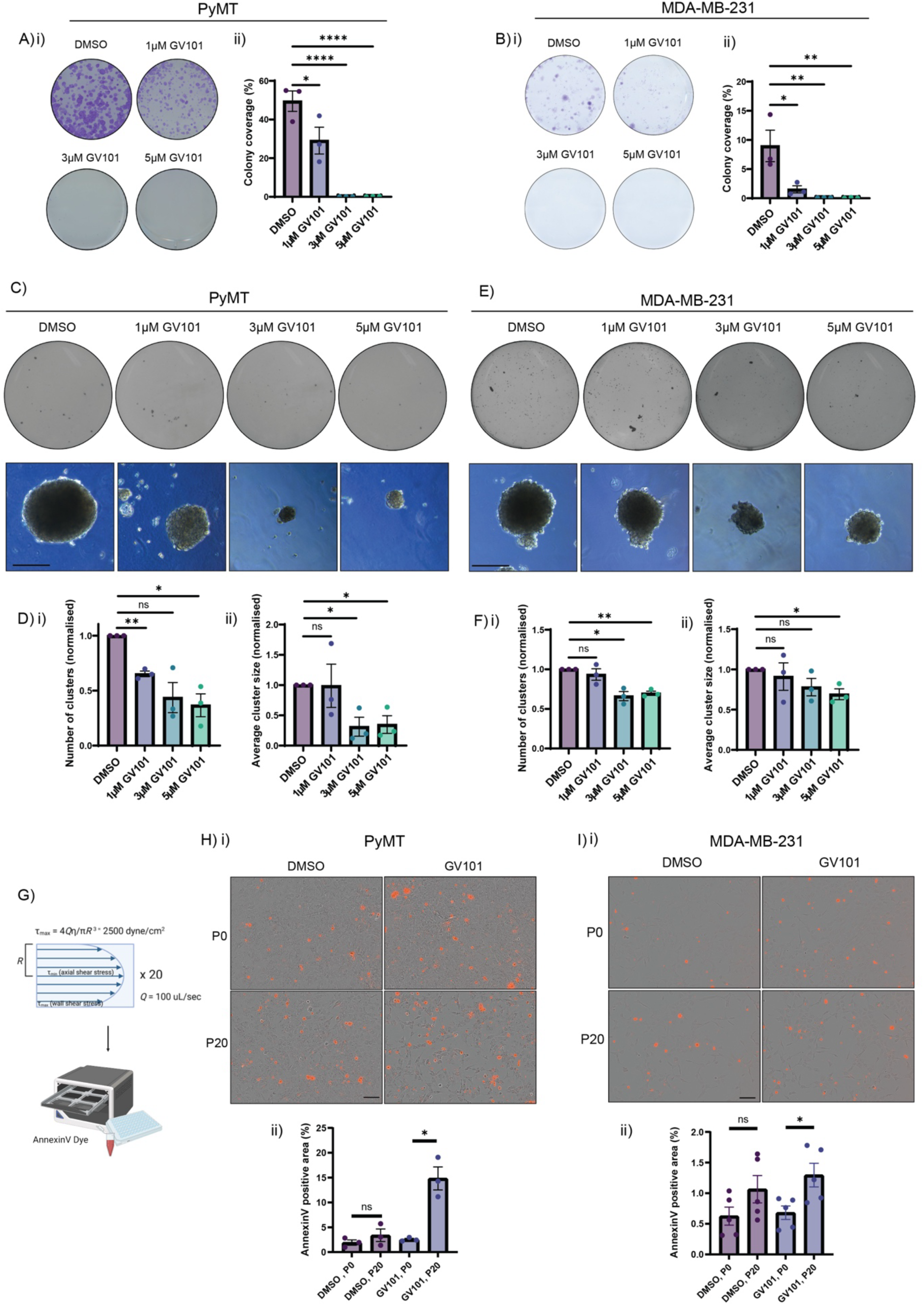
ROCK2 inhibition with GV101 reduces epithelial cell outgrowth in 2D and 3D while also sensitising cells to fluid flow-induced shear stress, also see Figures S2 and S3. **(A)** Representative images of PyMT cancer cell 2D colony formation assay stained with crystal violet to visualise colonies following 4 days of treatment (i) and quantification of colonies (ii). **(B)** Representative images of MDA-MB-231 2D colony formation assays stained with crystal violet to visualise colonies following 4 days of treatment (i) and quantification of colonies (ii). **(C)** Representative images of whole wells stained with quick-dip II to visualise PyMT cancer cell AIG clusters (top panel) and representative images of individual AIG clusters (bottom panel). **(D)** Quantification of PyMT cancer cell AIG cluster number (i) and average cluster size (ii). **(E)** Representative images of whole wells stained with quick-dip II to visualise MDA-MB-231 AIG clusters (top panel) and representative images of individual AIG clusters (bottom panel). **(F)** Quantification of MDA-MB-231 cancer cell AIG cluster number (i) and average cluster size (ii). **(G**) Schematic of shear stress assay to assess GV101 treatment following fluid flow-induced shear stress using AnnexinV incucyte imaging. **(H)** Representative images of PyMT cancer cells after fluid flow-induced shear stress (P0 = no shear stress, P20 = 20 cycles of shear stress) and subsequent treatment with DMSO or GV101 and AnnexinV dye (red) to visualise cell apoptosis (i) and quantification of PyMT cancer cell apoptosis at 24 hours after fluid flow-induced shear stress (ii). **(I)** Representative images of MDA-MB-231 cancer cells after fluid flow-induced shear stress (P0 = no shear stress, P20 = 20 cycles of shear stress) and subsequent treatment with DMSO or GV101 and AnnexinV dye (red) to visualise cell apoptosis (i) and quantification of MDA-MB-231 cancer cell apoptosis at 24 hours after fluid flow-induced shear stress (ii). Data represents mean ± SEM of three individual repeats. p-values were determined using One-sample t test for normalised data and Unpaired t-test for remaining data was used. For shear stress analysis an unpaired t test was used. ns = not significant (p≥0.05), * = p<0.05, ** = p<0.01, **** = p<0.0001. Scale bar = 150µm.

In addition, biomechanical stress is well-known to induce ROCK signalling in cells [20–22], which can protect cells from mechanical damage and subsequent cell death. We therefore next assessed the effects of GV101-mediated ROCK2 inhibition on cell survival following fluid flow-induced shear stress [35, 65, 70]. Here, we subjected cells to shear stress at a rate of 100 µL/s, as previously achieved (Figure 3G) [35, 65, 70]. Following 20 cycles of shear stress, single cells were seeded to observe colony outgrowth in the presence of GV101 or vehicle. Cell survival was assessed *via* live AnnexinV imaging of apoptosis followed by segmentation of cells and AnnexinV signal to determine the proportion of apoptotic cells (Figure S3A and Figure S3B). In PyMT cancer cells, shear stress alone did not increase apoptosis (Figure 3H, compare DMSO P0 and P20), while the addition of GV101 after shear stress exposure (P20) significantly increased apoptosis compared to vehicle (Figure 3H). Similarly, MDA-MB-231 cells showed a non-significant increase in apoptosis following shear stress which again increased with GV101 addition (Figure 3I), in line with the PyMT data. Overall, this suggests that ROCK2 is important for cell survival following fluid flow-induced shear stress and that ROCK2 inhibition with GV101 can expose cell vulnerability to apoptosis in this setting [71].

### ROCK2 inhibition reduces single cell migratory dynamics

Another well characterised role of ROCK signalling is its function as a driver of cell migration through regulation of the actin cytoskeleton and integration of cell-ECM or environment interactions [14]. This has been demonstrated during developmental processes [72, 73] as well as in disease contexts such as cancer, where co-opted ROCK signalling can promote cancer cell migration, invasion and metastasis [12, 13, 35]. To assess the effects of GV101 on single-cell migration across an ECM substrate, we first generated CDMs (Figure 4A) as described previously [35, 63, 66, 68, 69]. Following 7 days of matrix deposition, CDMs were de-cellularised leaving the native ECM behind, onto which cancer cells were then seeded (Figure 4A). Once adhered to the CDM, cells were treated with GV101 or vehicle and single cell migration was tracked longitudinally over a 12-hour period, as previously achieved [65, 67, 68]. As PyMT cancer cells were found to not be highly migratory on a CDM (data not shown), single-cell migration upon ROCK2 inhibition was assessed using the MDA-MB-231 cell line. Polar plots of individual cell tracks over the 12-hour period showed that GV101 treatment profoundly impaired the ability of cells to migrate on a CDM compared to vehicle (Figure 4B). Quantification of key cell migration readouts further confirmed a significant decrease in average distance travelled, average track displacement and maximum distance travelled over a 12-hour period for cells treated with GV101 compared to vehicle (Figure 4Ci to iii, Movie S7). In addition, a significant decrease in the migratory speed of these cells was also observed with GV101 treatment compared to vehicle (Figure 4Civ). Overall, this data demonstrates that ROCK2 is required for key locomotive processes of individual cells on a CDM.

**Figure 4:**
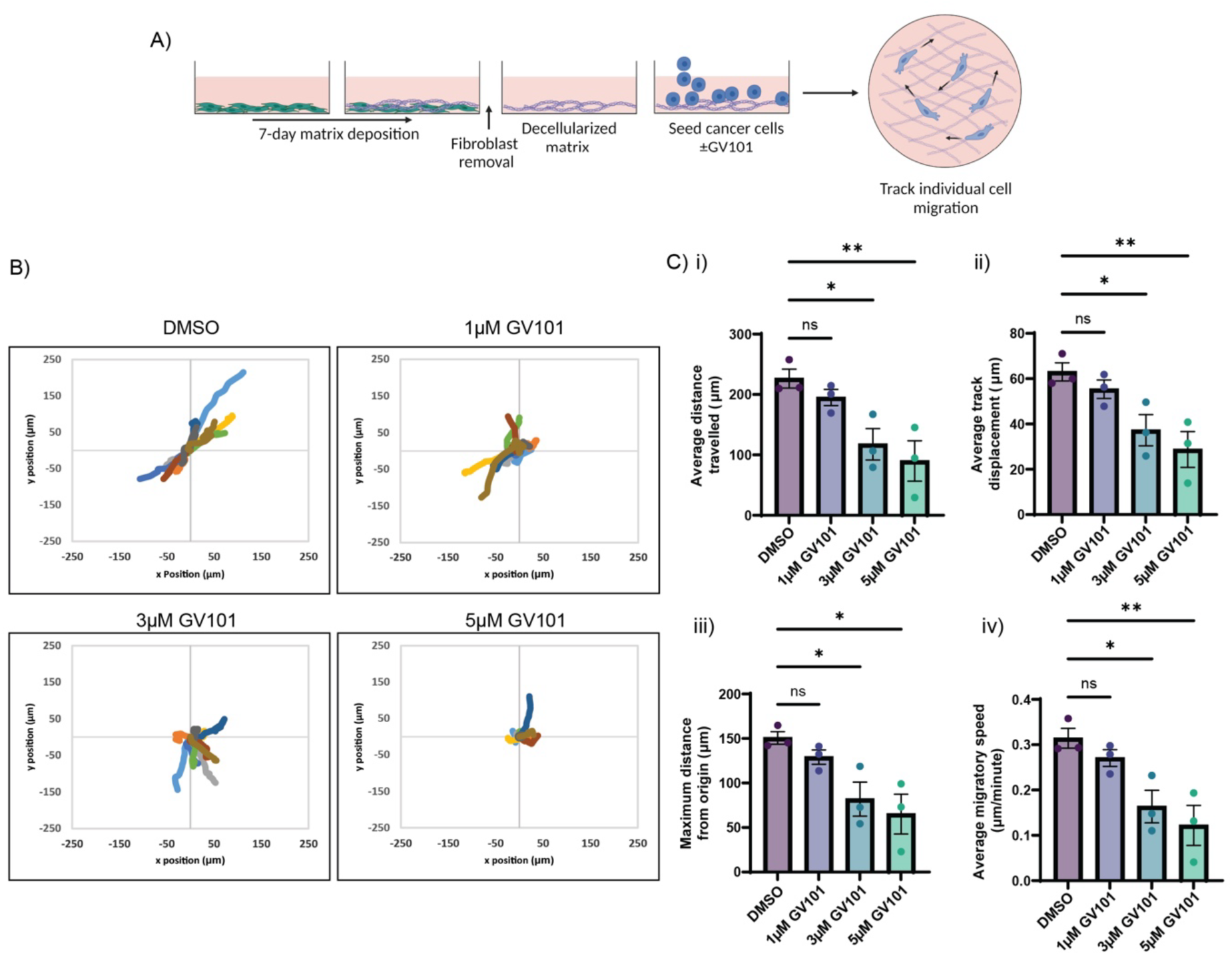
ROCK2 inhibition with GV101 decreases the single-cell migratory capacity of epithelial cells on cell-derived matrices. **(A)** Schematic of single-cell migration assay where MDA-MB-231 cells were seeded on cell-derived matrices (CDMs), treated with GV101 and individual migratory patterns tracked over a 12-hour period. **(B)** Representative polar plots showing cell migratory tracks of individual cells (depicted as individually coloured tracks) from a starting point (origin) over 12 hours of migration. **(C)** Quantification of single cell migration showing average distance travelled (i), average track displacement (ii), maximum distance from origin (iii) and average migratory speed (iv). Data represents mean ± SEM of three individual repeats, with 40 cells/condition/repeat analysed. p-values were determined using a One-way ANOVA with multiple comparisons, ns = not significant (p≥0.05), * = p<0.05, ** = p<0.01.

### Stromal ROCK2 governs subsequent collective migration dynamics of epithelial cells

While our initial studies focussed on understanding the requirement for ROCK2 in isolated fibroblast and epithelial cells, both cell populations co-exist *in vivo* where interactions between cancer cells and the adjacent ECM create a reciprocal activation of ROCK signalling. Moreover, in addition to single cell migration, cells can also move *via* collective migration, where the ECM can act as a guide to organise cellular migration [74–77]. This process is reliant on both intracellular cell migratory signalling as well as cell-cell and cell-ECM adhesions, all of which have been shown to be regulated by ROCK signalling. To investigate the effects of ROCK2 inhibition with GV101 on collective cell migration and cell-ECM interactions, migration assays were performed on CDMs with the aim of dissecting the contribution of stromal *versus* epithelial ROCK2 inhibition on collective cell migration (Figure 5Ai) [65, 68, 77]. Here PyMT cancer cells were utilised as MDA-MB-231 cells did not form highly organised collective migration phenotypes (data not shown). Firstly, the effects of ROCK2 inhibition *via* GV101 on cells to initiate cell-ECM contacts with the CDM after cell attachment was investigated. Here, we quantified circularity as a surrogate for cell elongation and attachment to the CDM 6 hours post seeding (Figure 5Aii and Movie S8). Cells treated with GV101 exhibited an increased circularity compared to vehicle (Figure 5Bi and ii), indicating a reduced capability to engage effective cell-ECM attachment upon ROCK2 inhibition. Collective cell migration was then assessed *via* measurement of cell streaming anisotropy following GV101 treatment, where a high anisotropy indicates highly organised migration whereas a low anisotropy correlates with disorganised migration (Figure 5C) [35, 65, 66, 78]. Here we observed a subtle decrease in organised cell migration when cells were treated during the migration phase with GV101 compared to vehicle control (Figure 5Di and ii and Movie S8). Overall, these data indicate that epithelial ROCK2 inhibition with GV101 significantly affects cancer cell attachment while only subtly reducing the organisation of collective cell migration.

**Figure 5:**
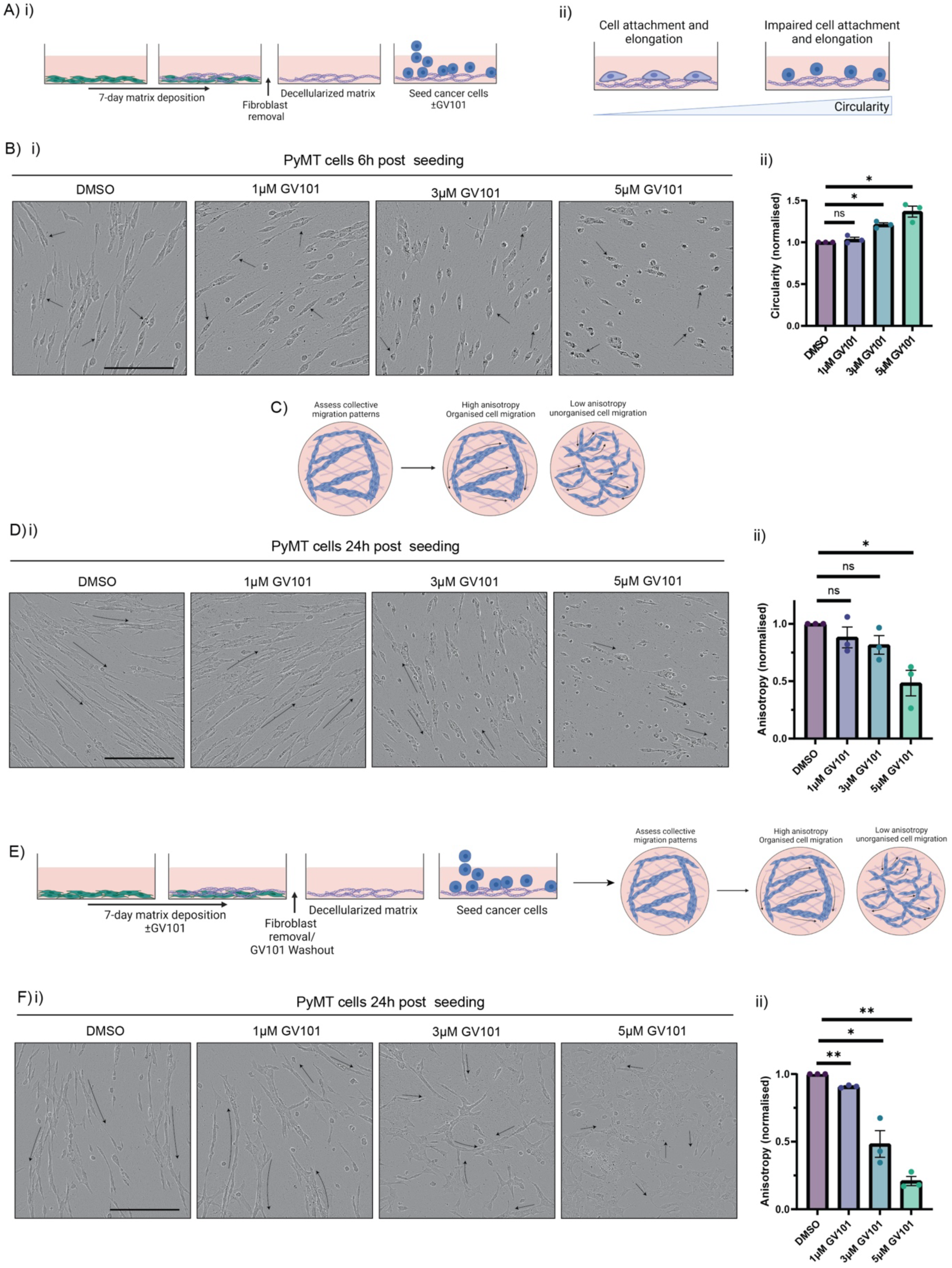
GV101 treatment of stromal ROCK2 reduces collective cell migration. **(A)** Schematic of cell-derived matrix (CDM) generation assay (i) and schematic of cell attachment and elongation on CDM to assess circularity (ii). **(B)** Representative images of PyMT cancer cell landing on a CDM following 6 hours of treatment with DMSO or GV101 (i) and quantification of PyMT cell circularity upon 6 hours of treatment (ii). **(C)** Schematic of collective cell migration showing organised (high anisotropy) and unorganised (low anisotropy) cell migration. **(D)** Representative images of PyMT cancer cell collective migration on a CDM following 24 hours of treatment with DMSO or GV101 (i) and quantification of PyMT collective cell migration *via* anisotropy (ii). **(E)** Schematic of CDM generation and PyMT cancer cell streaming assay upon GV101 treatment of fibroblasts during matrix deposition and assessment of cell streaming *via* anisotropy. **(F)** Representative images of PyMT cancer cells 24 hours post seeding on a GV101 pre-treated matrix (i) and quantification of collective cell migration *via* anisotropy (ii). Data represents mean ± SEM of three individual repeats. Data represents mean ± SEM of three individual repeats. p-values were determined using One-sample t test, ns = not significant (p≥0.05), * = p<0.05, ** = p<0.01. Scale bar = 200µm.

It is well appreciated that in addition to intracellular signalling driving cell migration, extracellular cues from the environment can affect cancer cell migratory behaviour. For example, subtle changes in ECM can be sufficient to disrupt co-ordinated cell migration [35, 65, 79, 80]. Given that we demonstrated a reduction in *de novo* ECM deposition and remodelling in CDM assays upon ROCK2 inhibition (Figure 2), we next assessed how GV101 primed CDMs, where GV101 was provided during the matrix production phase, would affect the subsequent ability of epithelial cells to undergo collective migration. Here, fibroblasts were primed with GV101 or vehicle during the CDM matrix deposition phase followed by de-cellularisation to remove the fibroblasts and subsequent GV101 washout to assess PyMT cancer cell streaming in the absence of GV101 (Figure 5E). Cells were allowed to grow on the CDMs for 24 hours prior to assessing collective cell migration *via* anisotropy quantification. Here, we observed a significant decrease in the organisation of cellular streaming on GV101 primed CDMs compared to vehicle (Figure 5Fi and ii, Movie S9). Interestingly, this decrease in organised collective cell migration on GV101 primed CDMs was more pronounced than when GV101 was provided to inhibit epithelial ROCK2 during the migration phase, highlighting a dominant role for the ECM substrate in guiding collective cell migration. Overall, this data demonstrates that while epithelial ROCK2 inhibition with GV101 perturbs cancer cell-ECM attachment, collective cell migration dynamics is predominantly disrupted when *de novo* ECM deposition and remodelling is altered with GV101 treatment of ECM-producing fibroblasts.

### Uncoupling epithelial *versus* stromal roles of ROCK2 during cancer cell invasion into 3D organotypic matrices

To further dissect the role of ROCK2 in the stromal *versus* epithelial compartment, we next used a 3D organotypic invasion assay, where cancer cells interact with and directionally invade into a fibroblast-contracted collagen matrix. Here, fibroblasts were embedded into a collagen matrix and allowed to contract the matrix over 12 days as described previously (Figure 1A) [35, 60, 63–66, 81–83]. Following contraction, MDA-MB-231 cancer cells were seeded on top of the fibroblast-contracted collagen matrices (Figure 6A). After 3 days of cell attachment and growth, the 3D organotypic cultures were placed on a metal grid to create an air-liquid interface producing a chemotactic gradient towards which the cancer cells were allowed to invade for 14 days (Figure 6A) [62, 66, 70, 83, 84]. Here, we aimed to uncouple the effects of stromal ROCK2 targeting with GV101 on creating an invasion-permissive environment by treating the CAFs during the contraction phase (early priming treatment) *versus* the effects of GV101 on epithelial ROCK2 function during cancer cell invasion (late invasion treatment) or in a continuous manner (chronic treatment, Figure 6A) as previously achieved [35, 65].

**Figure 6:**
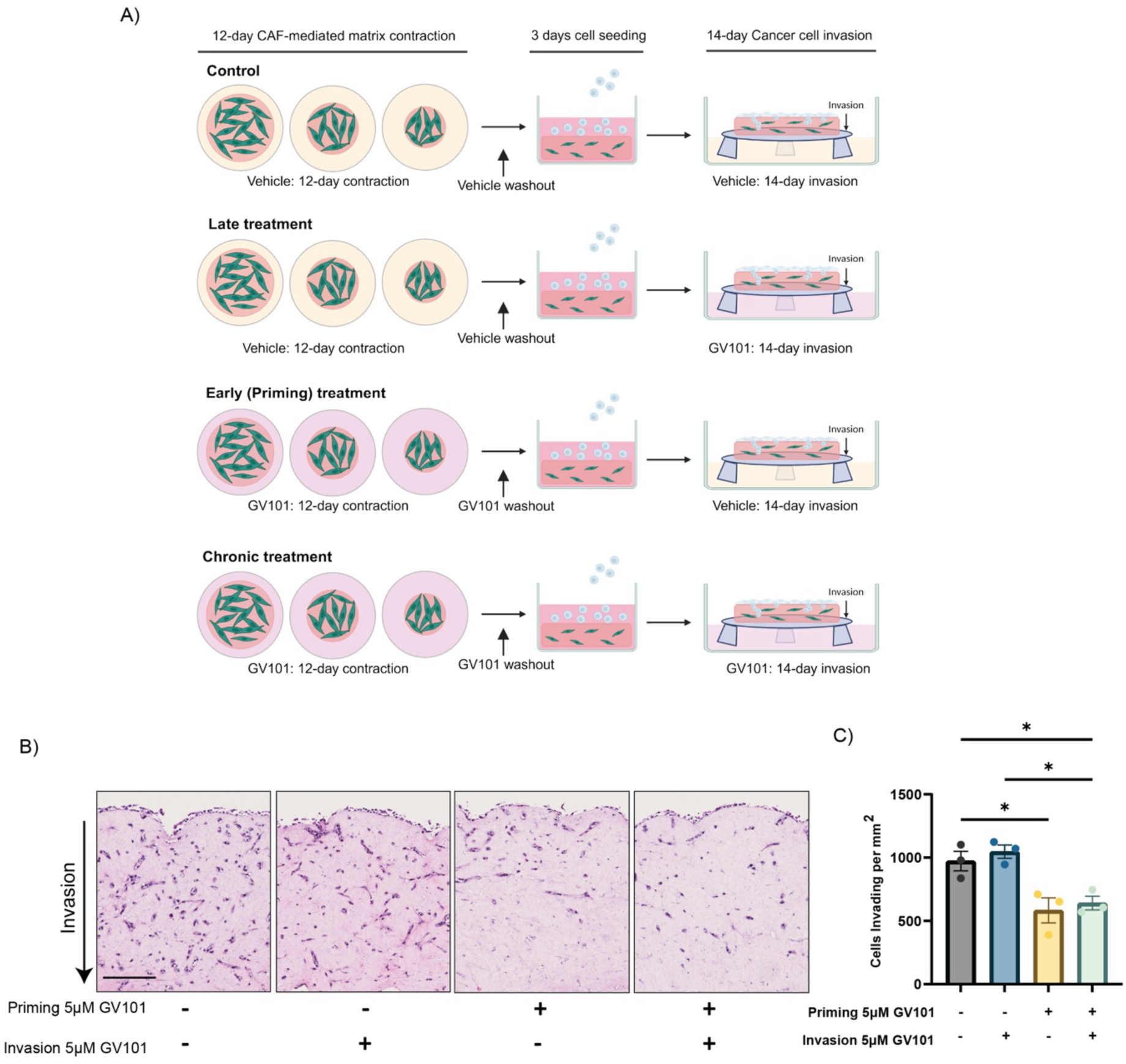
Disruption of stromal matrix remodelling impairs cell invasion into 3D organotypic collagen matrices. **(A)** Schematic of organotypic invasion assay showing fibroblast-mediated collagen contraction, cancer cell seeding and subsequent invasion into contracted matrices towards a chemotactic gradient. **(B)** Representative images of H&E-stained cross-sections of MDA-MB-231 cancer cell organotypic invasion assays with cancer cell invasion into the matrix assessed upon early stromal priming, late invasion treatment or continuous (chronic) treatment with GV101. **(C)** Quantification of number of MDA-MB-231 cancer cells invading into the matrix per mm^2^. Data represents mean ± SEM of three individual repeats. p-values were determined using a One-way ANOVA with multiple comparisons. * = p<0.05. Scale bar = 100µm.

Importantly, we observed that in the control setting MDA-MB-231 cells readily invaded into the collagen matrix (Figure 6B) and this was not significantly altered upon late invasion treatment *via* epithelial ROCK2 inhibition with GV101 compared to vehicle (Figure 6B and C). Interestingly, in the early priming treatment setting we observed a significant decrease in cancer cell invasion when we targeted stromal ROCK2 during matrix contraction (Figure 6B and C). Moreover, in line with previous studies [35, 65], continuous or chronic treatment by targeting both stromal and epithelial ROCK2 decreased cancer cell invasion to a similar level as early priming, suggesting that the disruption of stromal ECM remodelling *via* GV101 alone is sufficient to decrease subsequent cancer cell invasion. This data further supports our previous results on collective cell migration (Figure 5) where modulation of the ECM *via* stromal ROCK2 targeting with GV101 was sufficient to alter subsequent collective cell migration dynamics. Collectively, we uncouple how targeting epithelial and stromal ROCK2 govern cell movement with a role for epithelial ROCK2 in individual cell migration whereas stromal ROCK2 directs subsequent collective cell migration and invasion.

### ROCK2 expression increases during disease progression and correlates with poorer patient survival in TNBC cohorts

This study has demonstrated that ROCK2 inhibition with GV101 is able to reduce fibroblast-mediated remodelling of pre-existing and *de novo* synthesised collagen fibres. In addition, cell growth, proliferation and survival as well as cancer cell migration and invasion dynamics were also impaired upon GV101 treatment. Here we have utilised models of BC to systematically investigate the interactions between stromal and epithelial cells and their adjacent ECM due to the highly fibrotic, proliferative and invasive nature of this disease. ROCK inhibition has long been anticipated as a potential therapeutic target in a variety of cancer types, including BC, but efficacy and safety of previous generation ROCK inhibitors has hampered clinical translation [5]. The findings from this study suggest that ROCK2 inhibition may be able to impair cell proliferation, survival and invasion as well as reduce fibrosis in BC models. We therefore aimed to investigate the role of the ROCK2 isoform specifically in human BC patient cohorts. Analysis of both the METABRIC and TCGA BC cohorts revealed that high ROCK2 mRNA expression significantly correlated with poorer patient median survival, (Figure 7A and Bi). Interestingly, we observed in TCGA data that the level of ROCK2 mRNA expression was also significantly increased in the highly fibrotic and invasive TNBC subtype compared to other BC subtypes combined (Figure 7Bii), suggesting that ROCK2 could play a role in TNBC disease progression and survival.

**Figure 7:**
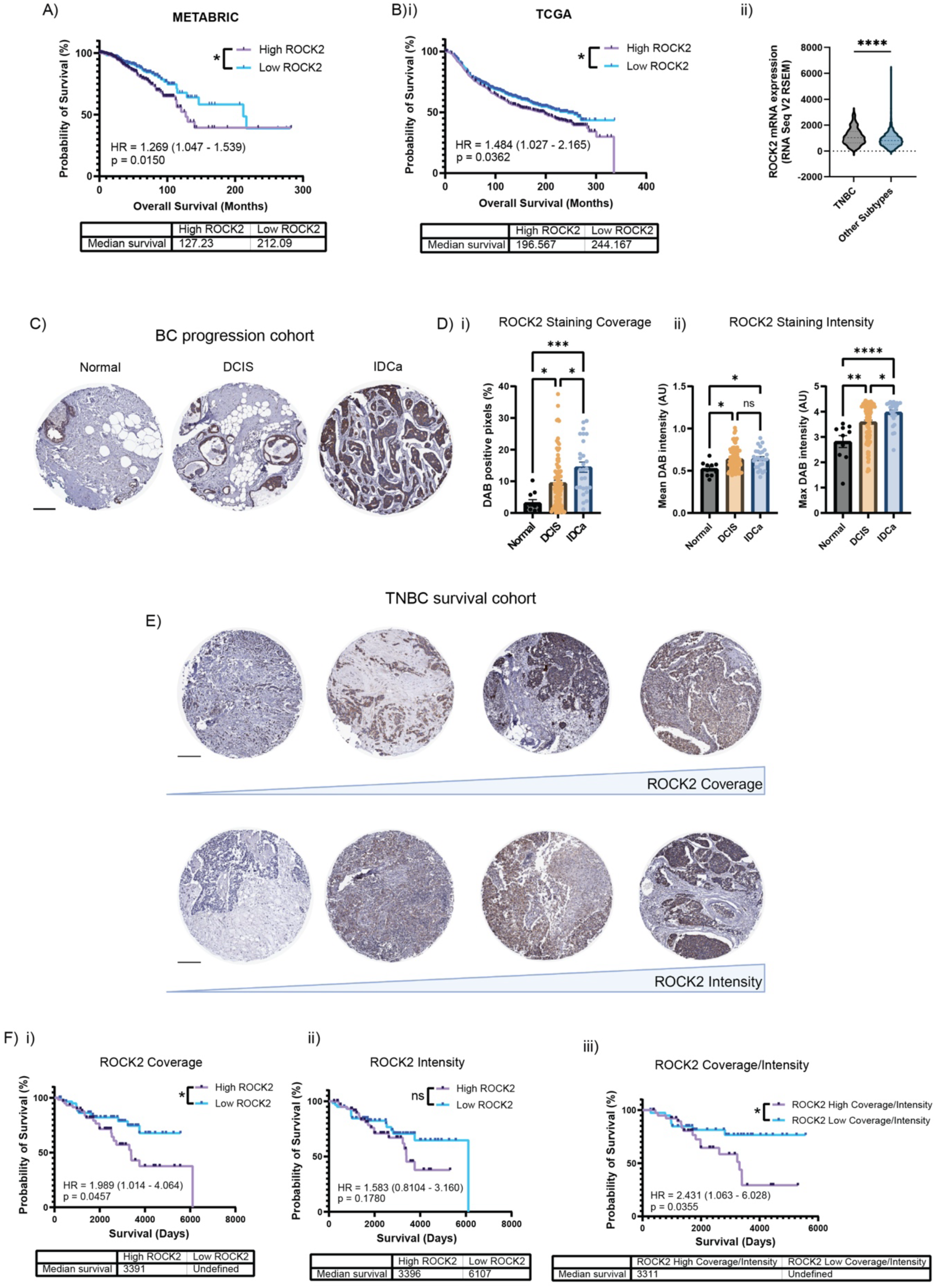
High ROCK2 expression correlates with disease progression and poorer survival in human breast cancer cohorts. **(A)** Survival analysis of breast cancer (BC) patients from the METABRIC BC patient cohort stratifying patients based on ROCK2 mRNA expression. **(B)** Analysis of the TCGA BC patient cohort assessing ROCK2 mRNA expression on patient survival (i) and comparing ROCK2 mRNA expression levels in triple-negative breast cancer (TNBC) patients to all other BC subtypes (ii). **(C)** Representative images of TMA cores from the Garvan/RPA Breast Progression Series stained with immunohistochemistry for ROCK2. **(D)** Quantification of ROCK2 staining coverage (i) and staining intensity (ii) in the Garvan/RPA Breast Progression Series TMA cohort. **(E)** Representative images of TMA cores stained with immunohistochemistry for ROCK2 from the RPA/CRGH TNBC TMA cohort showing increasing ROCK2 staining coverage and intensity. **(F)** Survival analysis assessing ROCK2 staining coverage (i), intensity (ii) and combined coverage and intensity (iii). For survival curves p-values were determined using Log-rank (Mantel-Cox) test and hazard rations (HR) with 95% confidence interval by a univariate Cox Proportional Hazards Model. For subtype comparison Unpaired t-test was used and for disease progression an Ordinary one-way ANOVA with multiple comparisons was used. * = p<0.05, ** = p<0.01, *** = p<0.001, **** = p<0.0001. Scale bar = 200µm.

We next interrogated ROCK2 protein expression *via* immunohistochemistry (IHC) in two independent human BC TMA cohorts. Firstly, the Garvan/RPA Breast Progression Series was used to assess ROCK2 protein expression over disease progression. Staining and analysis of the Garvan/RPA Breast Progression Series (Figure 7C) revealed that staining coverage was significantly increased at each stage of disease from healthy (normal) breast tissue to Ductal Carcinoma In Situ (DCIS) and again from DCIS to Invasive Ductal Carcinoma (IDCa, Figure 7Di). Mean and maximum staining intensity was also increased, from normal breast tissue to DCIS (Figure 7Dii). Furthermore, a significant increase in the average maximum intensity of the staining from normal tissue over DCIS to IDCa was observed (Figure 7Dii). Together this data shows that ROCK2 protein expression is significantly increased during BC disease progression.

Secondly the RPA/CRGH TNBC cohort was used to assess ROCK2 protein expression in relation to patient outcome in a TNBC-specific setting (Figure 7E). This revealed that high staining coverage significantly correlated with poorer patient survival (Figure 7Fi). Similarly, we observed a decrease in median survival for those patients with high staining intensity (Figure 7Fii). Moreover, combined assessment of the staining coverage and intensity scores showed a significant decrease in median survival for patients with high ROCK2 expression compared to those with low ROCK2 expression (Figure 7Fiii). This data supports the mRNA data and overall indicates that high ROCK2 expression in human TNBC patients correlates with poorer patient survival. Therefore, ROCK2 may represent a novel target in TNBC warranting further interrogation of ROCK2 specific inhibitors in this disease context.

## Discussion

The ROCK signalling axis has been well characterised in a variety of cellular processes including actomyosin contraction, cell proliferation, apoptosis and migration as well as tissue-level processes including ECM synthesis and remodelling [1, 6, 14]. As a result, dysregulation of this pathway has been implicated in multiple diseases including but not limited to autoimmune, cardiovascular and fibrotic diseases as well as solid cancers [5, 11, 24]. Inhibition of ROCK has been long suggested as a possible therapy for these diseases due to the multi-faceted roles of ROCK signalling. For example, in cancers that exhibit highly fibrotic signatures, such as pancreatic cancer and TNBC, ROCK inhibition was shown to act as a dual targeting strategy reducing stromal fibrosis and stiffness to improve therapy delivery and efficacy as well as directly limiting cancer cell proliferation and invasiveness [35, 56, 85]. These earlier studies have been limited to the use of pan-ROCK1/2 inhibitors, which often exhibit undesired side effects and toxicity. The generation of ROCK2 specific inhibitors with preferable safety profiles, such as Belumosudil, has enabled assessment of ROCK2 inhibition, which was successfully translated into humans for the treatment of chronic graft *versus* host disease [47, 86]. Most recently, GV101, which has 1000-fold more selectivity for ROCK2 over ROCK1, was reported to stall fibrosis in a mouse model of non-alcoholic fatty liver disease with increased efficacy over Belumosudil [49]. Here, we assessed the effects of GV101-mediated ROCK2 inhibition on both stromal and epithelial cell biology of BC. This study utilised TNBC as a model of a highly fibrotic, proliferative and invasive subtype of BC to investigate the effects of ROCK2 inhibition with GV101 on key cellular processes in both stromal and epithelial cell populations.

ECM synthesis and remodelling is often a defining feature of fibrotic diseases and has also been implicated in various solid cancers, such as TNBC. The ROCK1/2 signalling axis plays a crucial role in the remodelling and stiffening of the ECM through its control of actomyosin contraction [22, 36]. Here we show that ROCK2 inhibition with GV101 reduces fibroblast-mediated collagen remodelling of pre-existing collagen in organotypic matrices and of *de novo* synthesised collagen in CDM assays.

In addition to the regulation of ECM remodelling through actomyosin contraction, the ROCK signalling axis has also been shown to regulate other cytoskeleton-dependent processes including cell proliferation and survival [11, 87]; however, there is conflicting evidence as to the involvement of ROCK1/2 in these processes depending on cell type. In non-small cell lung carcinoma deletion of both ROCK1 and 2 was required to suppress cell proliferation [11], while in hepatocellular carcinoma and neuroblastoma decreased ROCK2 expression alone was sufficient to reduce proliferation [38, 39]. Here, we demonstrate that in the setting of TNBC, ROCK2 inhibition reduces baseline cell viability as well as colony outgrowth from individual cells in a 2D and 3D setting, again highlighting the context-dependent or disease-specific role of this signalling axis. Furthermore, upon cell detachment from the ECM, altered integrin signalling has been shown to activate ROCK2 resulting in cytoskeletal rearrangements preventing anoikic cell death, a key feature of cancer cells allowing them to survive in the absence of cell-substrate interaction under AIG conditions [16, 88]. In our study, inhibition of ROCK2 with GV101 under AIG conditions resulted in a significantly reduced ability of cancer cells to survive and grow out into colonies [35, 89]. Moreover, we also demonstrate that GV101 sensitises cells to apoptosis following biomechanical stress, which is known to induce cell response *via* multiple cytoskeletal rearrangements [71, 90, 91]. Overall, these data highlight a role for ROCK2 in cell proliferation and survival as well as in response to cell stressors such as AIG and fluid flow-induced shear stress conditions.

In this study we also aimed to uncouple the effects of both epithelial and stromal cell ROCK2 inhibition on subsequent cancer cell migration and directional invasion. We first investigated both single cell and collective cell migration dynamics on CDMs upon ROCK2 inhibition, which provide readouts of cell migratory capacity and also cell-ECM adhesion interactions. The ROCK signalling axis governs both processes through its activation by upstream kinases, such as RhoA and Focal Adhesion Kinase (FAK) from ECM-mediated input, as well as its downstream regulation of the actin cytoskeleton. In this study we demonstrated that ROCK2 inhibition with GV101 reduces both single cell migratory distance and speed. In the context of collective cell migration, epithelial ROCK2 inhibition with GV101 reduced initial cell-ECM attachment, elongation and spreading. Interestingly, collective migration dynamics on a CDM were only subtly altered following epithelial GV101 treatment, whereas manipulation of the matrix on which the cells were seeded, *via* GV101 inhibition in fibroblasts during the ECM deposition phase, resulted in a profound reduction of organised cell streaming. Importantly, this was superior to epithelial ROCK2 inhibition alone. In addition, we demonstrated that GV101 disrupted the deposition and remodelling of *de novo* synthesised ECM in a CDM setting and these results highlight that, in particular stromal ROCK2 inhibition, has the capacity to alter subsequent collective cell migration dynamics by modulating the extrinsic cell-ECM adhesions. Interestingly, manipulation of the stromal ECM environment has also been observed to affect collective cell migration in other contexts, for example FAK inhibition during CDM production affects collective pancreatic cancer cell migration predominantly through cell extrinsic mechanisms [65].

Our results from CDM assays were further corroborated in a 3D context of directional cell invasion into collagen matrices, where manipulation of fibroblast-mediated ECM remodelling *via* stromal ROCK2 inhibition with GV101 was also sufficient and superior over epithelial ROCK2 inhibition to reduce subsequent cancer cell invasion. Given the considerable effects that epithelial ROCK2 inhibition has on reducing both cell growth and cell migration, the lack of efficacy in reducing cell invasion during the late invasion treatment was surprising with only early stromal priming treatment reducing invasion. However, this has also been observed with pan-ROCK1/2 inhibition in other contexts. For example, in pancreatic cancer, pan-ROCK1/2 inhibition with Fasudil in the stromal compartment during fibroblast-mediated matrix remodelling was sufficient to impair subsequent cancer cell invasion into 3D organotypic invasion assays, to a greater extent than treatment of epithelial ROCK1/2 during the invasion phase [35]. Similar effects were observed upon FAK inhibition, where stromal FAK inhibition reduced epithelial cell invasion [65]. Furthermore, the biomechanical properties of the surrounding environment have been demonstrated to influence cancer cell invasion dynamics. Inhibition of lysyl oxidase (LOX), a collagen-crosslinking enzyme, during collagen matrix remodelling reduced matrix stiffness and subsequent cancer cell invasion [64]. In our study, we found that inhibiting stromal ROCK2 with GV101 during matrix contraction reduced the matrix stiffness and these biomechanical changes in the matrix are likely to contribute to the decreased cancer cell invasion observed. Overall, this work demonstrates the impact of cell-ECM outside-in signalling on cancer cell invasion and how manipulation of the surrounding ECM has effects on cell migratory and invasion dynamics, which could be utilised to reduce cell motility in highly invasive cancers, such as BC.

The ROCK signalling pathway has been implicated in a number of solid cancers where high ROCK2 mRNA and protein expression has been correlated with more aggressive disease and poorer patient survival [36, 92–96]. ROCK1/2 has also been found to be associated with BC metastasis with higher levels of ROCK1/2 mRNA found in metastatic sites when compared to matched primary tumours [97]. Most recently it has been shown that ROCK1/2 protein expression and phosphorylation of its downstream target, MYPT1, is increased in IDCa compared to normal/hyperplastic breast tissue [23] overall highlighting ROCK signalling as potential target. Here, we assessed ROCK2 specifically in relation to human BC disease showing that ROCK2 expression is increased during BC progression and that high ROCK2 expression correlates with poor patient outcomes in TNBC, the most lethal subtype of BC. Given the highly invasive and fibrotic nature of this disease, therapeutic interventions that target both epithelial and stromal cell populations, such as ROCK2 inhibitors may be a therapeutic option in this disease. However, given the highly context-dependent nature of ROCK2 signalling more studies are needed to investigate ROCK2 inhibitors in disease-specific settings.

Overall, this study has set out to examine and uncouple the effects of ROCK2 inhibition *via* GV101 on stromal and epithelial cellular function in isolated and co-culture models. Our findings demonstrate that inhibition of stromal ROCK2 with GV101 can reduce the ECM deposition and remodelling ability of stromal cells, while epithelial ROCK2 inhibition decreased cancer cell proliferation and survival. Furthermore, we have demonstrated that disrupting stromal ECM deposition and remodelling *via* stromal ROCK2 inhibition is sufficient to decrease subsequent collective cancer cell migration and invasion. Finally, we also reveal that high ROCK2 expression correlates with poor patient outcomes in the TNBC subtype of BC. Given these data we suggest that ROCK2 inhibitors should be further explored in the context of other fibrotic diseases and solid cancers. In line with this, ROCK2 inhibition may provide a dual targeting strategy to reduce both the fibrotic environment as well as decrease cancer cell proliferation, warranting further assessment of this signalling node in BC.

## Resource availability

### Lead contact

Further information and requests for resources and reagents should be directed to and will be fulfilled by the lead contact, David Herrmann.

### Materials availability

All materials generated in this study will be available upon reasonable request from the lead contact.

### Data and code availability

All information required to reanalyse the data reported in this study is available from the lead contact upon reasonable request.

## Acknowledgements

This study was supported by the following facilities at the Garvan Institute of Medical Research: Australian BioResource (ABR), Biological Testing Facility (BTF), Garvan Molecular Genetics (GMG), Tissue Culture Facility under R. Lyons, the Garvan Imaging Platform under D. Barkauskas, and the Garvan Histopathology and Biospecimen Facility under Anaiis Zaratzian. Some imaging was performed at the UNSW Katharina Gaus Light Microscopy Facility. We also would like to thank the staff of the ACRF INCITe Centre under co-leadership of T. G. Phan and P. Timpson. We would also like to thank our consumers (E. Jurd, V. Killen, J. Mumford and D. Goulburn OAM) for their valuable input regarding this work. Schematics were created with BioRender.com.

This study was supported by the National Health and Medical Research Council (NHMRC: 1136974, 1158590, 1160022, 1188208, 2000937, 2010330, 2012837, 2016930, 2018440, 2028766), US Department of Defense Breast Cancer Research Program (BC210277), Cancer Council NSW (RG 19-09, RG 21-12, RG 23-07 and RG 23-11), Cancer Institute NSW (CINSW; ECF012, ECF011, ECF1309, ECF1384, ECF1006, CDF1071 and TPG2100), Tour de Cure, St. Vincent’s Clinic Foundation, an Australian Cancer Research Foundation (ACRF) infrastructure grant (INCITe Centre), Estée Lauder Award and Suttons family and Len Ainsworth foundation philanthropy. P.T. was supported by a National Health and Medical Research Council (NHMRC) Fellowship (1136974 and 2016930) and a Len Ainsworth Fellowship. T.R.C. was supported by an NHMRC Investigator award (2033065) and Career Development Fellowship (1158590) and CCNSW (24-06). K.W. was supported by an Australian Government Research Training Scholarship. A.S. (2018440) was supported by a NHMRC Investigator Fellowship. M.N. (ECF012), S.R. (ECF1006), K.J.M. (ECF1384), B.A.P. (ECF1309) and D.H. (ECF011) were supported by a CINSW Early Career Research Fellowship. J.Z. was supported by a UNSW SPHERE Cancer CAG PhD scholarship. M.T. was supported by a White Walker Cancer Research PhD scholarship. S.R. was supported by a UNSW International PhD scholarship. M.S.S. was supported by an Australian Breast Cancer Research Fellowship from The Hospital Research Foundation. D.A.R. and C.R.C. were supported by a Baxter Family Postgraduate scholarship and Australian Government Research Training Scholarship.

## Author contributions

**Investigation, validation, and formal analysis:** All authors. **Conceptualisation and funding acquisition:** D.A.R., M.N., D.G.-O., A.J.G., T.R.C., B.A.P., K.J.M., E.L., A.S., S.O’T., M.S.S., C.E.C., A.Z-Z., P.T. and D.H. **Writing, editing, and visualisation:** D.A.R., P.T. and D.H.

## Declaration of interests

P.T. and D.H. receive reagents from Équilibre Biopharmaceuticals and Graviton Biosciences (GV101 provided free of charge). P.T. receives reagents from Kadmon Inc., InxMed (consultant), Redx Pharma, and Amplia Therapeutics. Under a licensing agreement between Amplia Therapeutics and Garvan Institute of Medical Research, K.J.M., P.T. (consultant) and D.H. are entitled to milestone payments.

## STAR Methods

### Experimental models and study participant details

#### Statistical analysis

Unless otherwise stated, p-values were determined using an Unpaired t test for comparisons between two groups or an Ordinary One-way ANOVA with Tukey’s multiple comparison for comparisons between three or more groups. For normalised data a One sample t test was used. For survival analysis Kaplan-Meier survival curves were analysed using a log-rank (Mantel-Cox) test. All statistical analysis was performed using GraphPad Prism (GraphPad Software Inc., CA, USA) with statistical significance defined as ns = p ≥ 0.05, * = p < 0.05, ** = p < 0.01, *** = p < 0.001, **** = p < 0.0001.

#### Human ethics

All research involving TMA biospecimens and associated data was covered by the Royal Prince Alfred Hospital Human Ethics Review Committee Approval (X14-0241) with all patient data supplied in a deidentified format.

#### Cell culture

PyMT 20065 cancer cells, herein referred to as PyMT cancer cells, previously generated by Dr Karen Blyth at the CRUK Beatson Institute (Glasgow, UK) [70], were maintained in DMEM (Dulbecco’s modified Eagle medium, Gibco) supplemented with 10% Fetal Bovine Serum (FBS), 1% penicillin/streptomycin (P/S), 5µg/mL insulin, 10ng/mL epidermal growth factor (EGF) and 10ng/mL Cholera Toxin A. PyMT CAFs, previously isolated from end-stage MMTV-PyMT tumours [37] were kindly provided by the lab of Dr Fernando Calvo lab and were maintained in DMEM supplemented with 10% FBS, 1% P/S, and 1x Insulin-Transferrin-Selenium (ITS-G, Gibco). MDA-MB-231 cells and Telomerase-immortalised fibroblasts (TIFs) [62] were maintained in DMEM supplemented with 10% FBS and 1% P/S. Cells were maintained at 37°C and 5% CO_2_ in a HeraCell 160i CO_2_/O_2_ incubator. Cell lines were tested for mycoplasma and confirmed to be negative. Primary human TNBC patient CAFs, obtained from Professors Elgene Lim and Alex Swarbrick at the Garvan Institute of Medical Research, were maintained in alpha modified Eagle’s media (αMEM) supplemented with 20% FBS and 1x Antibiotic/Antimycotic (AB/AM, Gibco) in collagen coated flasks. For collagen coating, flasks were incubated in 2mg/mL rat tail extracted collagen at 37°C for 2 hours. Following incubation flasks were washed twice with phosphate-buffered saline (PBS) and then incubated in growth media for one hour. Cells were maintained at 37°C, 5% O_2_ and 5% CO_2_ in a HeraCell 160i CO_2_/O_2_ incubator. All *in vitro* experiments were carried out in the growth media stated above, unless stated otherwise.

### Method details

#### Collagen extraction

Collagen I was isolated from rat tails as previously described [62]. Briefly, frozen rat tails were thawed in 80% ethanol and collagen tendons removed from the tail using tweezers. Extracted tendons were solubilised in 1.5L of 0.5M acetic acid (Fisher Chemicals) at 4°C for 48 hours with constant stirring using a magnetic stirrer. Following solubilisation the mixture was filtered through a moist gauze to remove any insolubilised tendons or sheath. Collagen I was then precipitated with 10% (w/v) NaCl at 4°C using a magnetic stirrer until a homogenous solution was achieved (approximately 6-8 hours). The solution was then centrifuged at 10,000 RPM for 30 minutes at 4°C to isolate precipitated collagen. Pelleted collagen precipitate was then dissolved in 0.25M acetic acid at a 1:1 ratio for 24 hours at 4°C using a magnetic stirrer. The collagen I solution was then dialysed in 17.4mM acetic acid for 3 days with acetic acid solution being replaced every 12 hours. Following dialysis the mixture was centrifuged at 14,000 RPM at 4°C for 90 minutes to remove any debris. Collagen concentration was measured using the Sircol™ Soluble Collagen Assay Kit (Biocolour) and was made up to a final solution of 2mg/mL in 17.4mM acetic acid prior to use.

#### Organotypic contraction assay

Organotypic contraction assays were performed as previously described [35, 60, 62, 64–66]. Rat tail extracted collagen I dissolved in acetic acid at a concentration of 2 mg/mL was neutralised with 0.22M sodium hydroxide with MEM 10x (Gibco). The neutralised collagen was kept on ice to prevent polymerisation. CAFs were prepared in FBS and then added to neutralised collagen I solution, 2.5mL of CAF/Collagen mixture was added to each well of a tissue culture (TC) treated plastic 6-well plate (Corning) for a final concentration of 3.5×10^5^ PyMT CAFs or 2.5×10^4^ TNBC human CAFs per well. Collagen/CAF matrices were placed at 37°C, 5% CO_2_ until solidified. The solidified collagen matrices were detached from the plate using a sterilised glass stripette, following addition of 5mL of appropriate growth media containing 5µM GV101, 5µM Fasudil or DMSO control. Matrix contraction took place over a 12-day period with drug containing media being replaced every 3 days along with the scanning of plates using a HP Flatbed Scanner (Hewlett Packard) to assess matrix size during contraction. Following 12 days of contraction matrices were fixed in 10% neutral buffered formalin for histopathology processing. Matrix size was quantified using ImageJ (National Institute of Health (NIH), USA) with three matrices analysed per condition for three independent repeats.

#### Unconfined compression analysis

Unconfined compression analysis was performed on organotypic matrices using a TA Instruments Dynamic Hybrid Rheometer (DHR-3, TA Instruments), in line with previously published work [64, 98]. Hole-punch biopsies with a diameter of 8mm were taken from organotypic collagen contraction matrices following 12 days of CAF mediated collagen contraction. Biopsies were placed between 80mm parallel plate geometries and a constant linear compression rate of 10µm/s was applied with axial force and gap width being measured every one second. Data was analysed by creating a stress/strain curve from axial force and gap measurements and the Young’s modulus was determined from the linear region of the stress/strain curve with three matrices analysed per condition for three independent repeats.

#### Cell-derived matrix deposition assay

Cell-derived matrices (CDMs) were generated as previously published [68, 69]. PyMT CAFs (6×10^4^ cells/well) or human TNBC CAFs (2×10^5^ cells/well) were seeded in a TC treated plastic 24-well plate. After 24 hours, confluent monolayers of CAFs were treated with 1mL of media containing 50µg ascorbic acid (Sigma) to stimulate matrix production. Fresh ascorbic acid was added directly to wells on days two and six with a complete media, drug and ascorbic acid refresh on day 4. Following one week of matrix production matrices were fixed in 10% neutral buffered formalin for 24 hours followed by dehydration in 70% ethanol for 24 hours before being imaged for SHG and subsequent picrosirius red staining.

#### Cell viability assay

PyMT 20065 cells (5×10^2^ cells/well) or MDA-MB-231 cells (4×10^4^ cells/well) were seeded in TC treated plastic 96-well plates (Corning) in appropriate growth media. Following 16 hours of adhesion cells were treated with 1, 3, 5µM GV101, 5µM Fasudil or DMSO control. Following 5 days of treatment cell viability was read out using an AlamarBlue™ Cell Viability Reagent Kit (ThermoFisher). Briefly 10uL of AlamarBlue reagent was added to wells containing 200µL of media and absorbance was read out on a FLUOstar® Omega Microplate reader (BMG Labtech) at an excitation of 544nm and emission of 590nm following 3.5 hours (PyMT 20065) or 5 hours (MDA-MB-231) of incubation. Cell viability was assessed in five wells per condition for three independent repeats.

#### Colony formation assay

Cells were trypsinised with 0.5% Trypsin-EDTA (Gibco) then filtered sequentially through 100µm, 70µm and 40µm filters to obtain single cell suspensions. Cells were seeded at 500 cells/well (PyMT 20065) or 1.0×10^3^ cells/well (MDA-MB-231) in TC treated plastic 6-well plates (Corning). Following cell attachment for 16 hours, cells were treated with 1, 3 or 5µM GV101 or DMSO control in appropriate growth media. For PyMT cells, cells were fixed on day 4 and for MDA-MB-231 cells were fixed on Day 14 by placing plates on ice and fixing with ice-cold 100% methanol for 20 minutes. Following fixation wells were washed twice with PBS (Gibco) followed by staining with 0.5% crystal violet (Merck) solution for 4 hours. Following staining wells were washed in water until excess crystal violet stain was removed and then dried upside down overnight. Colony coverage was assessed using ImageJ (NIH) in three wells per condition for three independent repeats.

#### Anchorage-Independent Growth Assays

AIG assays were performed from a modified published protocol [65, 70]. Preparations of 10% and 3% (w/v) low melting point agarose (SeaPlaque Agarose, Lonza) in H_2_O was autoclaved at least 24 hours prior to assay commencement. On the day of set-up both concentrations of agarose were microwaved briefly and warmed in a water bath at 42°C until molten. The 1% agarose solution was prepared by diluting the autoclaved 10% solution in the appropriate phenol red free media. A bottom layer of 1% agarose was made by adding 500µL of 1% agarose to the bottom of a TC treated plastic 24-well plate and allowed to solidify at room temperature for one hour. Cells were trypsinised with 0.5% Trypsin-EDTA (Gibco) then filtered sequentially through 100µm, 70µm and 40µm filters to obtain single cell suspensions. The 0.3% molten agarose was combined at a 2:1 ratio with media containing resuspended single cell solutions and 750µL of that final solution was added to 1% agarose base layer for a final concentration of 1.0×10^4^ cells/well (PyMT 20065) or 5.0×10^3^ cells/well (MDA-MB-231). Following 45 minutes incubation at room temperature 750µL of the appropriate phenol red free media was added with GV101 or DMSO control for final concentrations of 1, 3 and 5µM GV101. PyMT 20065 cells were grown in AIG conditions for 14 days and MDA-MB-231 cells for 10 days. Following completion of the assay individual AIG clusters were imaged using a DM4000 Leica microscope (Leica Biosystems, 10 clusters/well) to assess average cluster size. To assess total number of clusters 10µL of QuickDip II dye (Fronine Laboratory Supplies) was added to each well and following 24 hours of incubation whole well pictures were acquired using a Leica MZ12.5 microscope (Leica Biosystems). Average colony size was quantified in QuPath [99] via manual annotation and colony number quantified in ImageJ. AIG colony formation was assessed in four wells per condition for three individual repeats.

#### Flow-induced Shear Stress Assays

Flow-induced shear stress was performed as previously described [35, 65, 100]. PyMT or MDA-MB-231 cancer cells were trypsinised and filtered through a 70µm filter to a single cell suspension at a concentration of 5×10^5^ cells/mL in appropriate growth media. Cells were exposed to 20 rounds of shear stress through a 30-gauge needle at a constant flow rate of 100µl/s. After exposure to shear stress 2×10^3^ cells were seeded in a 96 well plate with Incucyte® Annexin V Red dye (Sartorius) and imaged on the Incucyte S3 live cell analysis instrument (Sartorius). For analysis cells were segmented using the Cyto3 model in Cellpose and mask created to quantify cell coverage in ImageJ. Masks of Annexin V were also generated in ImageJ and Annexin positive area was calculate from a ratio of Annexin V positive area divided by total cell coverage, with 9 ROIs per well for 3 wells analysed per condition for three independent repeats.

#### Single-Cell Migration Assays

Single-cell migration assays were performed from a modified protocol as previously published [65]. CDMs were generated as above with Telomerase-immortalised fibroblasts (TIFs). Following matrix production TIF-derived matrices were denuded via incubation in extraction buffer (20nm NH_4_OH, 0.5% Triton X-100, 1% Sodium Deoxycholate in PBS) for 3 minutes. Following cell removal matrices were carefully washed twice in PBS containing calcium and magnesium (1mM CaCl_2_ and 1mM MgSO_4_) and incubated in 10ug/mL DNase I (Roche) in PBS containing calcium and magnesium at 37°C, 5% CO_2_ for 30 minutes. Matrices were then washed twice in PBS containing calcium and magnesium before incubation in appropriate cell culture medium. MDA-MB-231 cells were trypsinised with 0.5% Trypsin-EDTA (Gibco) then filtered sequentially through 100µm, 70µm and 40µm filters to obtain single cell suspensions and seeded at 1.0×10^5^ cells/well on a TC-treated plastic 24-well plate following CDM production and denuding. Following cell attachment and elongation to matrix (6 hours) media was gently removed and replaced with fresh media containing 1, 3, 5µM GV101 or DMSO control. Wells were imaged every 30 minutes for 12 hours on an IncuCyte S3 Live-Cell Analysis Instrument (Sartorius). Single cell migration was then analysed using CellTracker image processing software [101], with 10 cells per well tracked for four wells per condition for three independent repeats (total 40 cells/condition/repeat).

#### Collective Cell Migration Assays

Collective cell migration was performed as previously reported [35, 65, 76, 77]. PyMT 20065 cells were trypsinised with 0.5% Trypsin-EDTA (Gibco). Cells were seeded onto denuded TIF derived CDMs at 5.0×10^5^ cells/well. Cells were seeded in media containing 1, 3, 5µM GV101 or DMSO control and imaged at 6 and 24 hours post seeding on an IncuCyte S3 Live-Cell Analysis Instrument (Sartorius). Cell attachment and elongation were assessed via the circularity function in ImageJ at 6 hours post seeding. Growth of cells on CDM and collective cell migration was assessed 24 hours post cell seeding using the ImageJ plugin FibrilTool [78]. Briefly, masks of cells were made from images and images were separated into quadrants before quantification of anisotropy (used as a surrogate readout for collective cell migration) using FibrilTool [78]. In total, nine regions of interest (ROI) were analysed per well with four wells per condition over three independent repeats.

#### Organotypic Invasion Assays

Organotypic invasion assays were performed using a modified version of previously published work [35, 60, 62, 65]. Matrices containing 1.67×10^5^ TIFs were treated with 5µM GV101 or DMSO control during contraction, with media and drug changed on days 3, 6 and 9. Following 12 days of contraction fibroblasts were killed with 400µg/mL Hygromycin B (Invitrogen) in growth media for 3 days. Matrices were then transported into individual wells of a TC-treated plastic 24-well plate. MDA-MB-231 cells (1×10^5^ cells/matrix) were seeded on top of TIF contracted matrices in appropriate growth media. Cells were allowed to grow on top of matrices for 3 days, during which a confluent monolayer was established. Matrices were then transferred onto metal grids which were placed in a 6cm TC treated plastic dish. Growth media containing 5µM GV101 or DMSO control was added to dishes to the level of the metal grid to create an air-liquid interface generating a chemotactic gradient, allowing cancer cells to invade into the collagen matrix. Media was changed every two days for a period of 14 days. Following 14 days of invasion matrices were fixed in 10% formalin for further histopathology processing and H&E staining. Analysis was performed using Qupath [99], with three matrices analysed per condition for three independent repeats.

#### Tissue processing, embedding and microtomy

All histopathology was performed in-house at the Histopathology and Biospecimen Facility at the Garvan Institute of Medical Research. Organotypic matrices were fixed in 10% neutral buffered formalin for 24 hours then transferred into 70% Ethanol for tissue processing using the Leica Peloris II (Leica Biosystems). During processing samples were moved through a graded series of Ethanol concentrations, followed by xylene, then paraffin at 60°C. Samples were then embedded using the TissueTek Embedding Station (Sakura Finetek).

Paraffin embedded blocks were sectioned at a thickness of 4µm using a Leica Microtome RM2235 (Leica Biosystems). Sections were placed onto Superfrost™ plus slides (Thermo Fisher Scientific) and allowed to incubate for 2 hours in a 60°C oven, for maximum adhesion.

#### Immunohistochemistry

All IHC staining was performed on a Leica Bond RX platform (Leica Biosystems). Slides were dewaxed (Bond Dewax Solution Ref: AR9222), followed by Heat-Induced Epitope Retrieval (HIER) using pH9 Epitope Retrieval solution 2 (AR9640, Leica Biosystems) for 20-30 minutes at 93°C for organotypic matrices or 100°C for tissue. Slides were incubated with primary antibody for 60 minutes, followed by a secondary antibody (polymer), then visualised with DAB (Bond Polymer Refine Detection Kit DS9800). All IHC slides were counterstained with Haematoxylin (Australian Biostain Haematoxylin Harris non-toxic, acidified) and coverslipped.

#### Picrosirius Red Staining

Picrosirius red staining was performed as achieved previously [35, 60, 65, 66]. Slides were dewaxed and counterstained with Haematoxylin (Australian Biostain Haematoxylin Harris non-toxic, acidified) for 4 seconds on a Leica ST5010 Autostainer XL (Leica Biosystems). Slides were then incubated in 0.02% phosphomolybdic acid for 2 minutes and briefly washed in water before a 2 hour incubation in 0.1% Picrosirius Red solution (Australian Biostain solution). This was followed by washes in acidified water prior to dehydration and clearing on a Leica ST5010 Autostainer XL (Leica Biosystems) and coverslipping. For CDMs, the CDMs were stained in wells using the same staining reagents mentioned above.

#### Imaging and Analysis

##### Picrosirius Red and Polarised Light Microscopy

Picrosirius red stained matrices were prepared as previously described and slides were scanned on the Olympus VS200 slide scanner (Olympus Life Science) at 20x magnification using both brightfield and polarised light imaging modalities. Picrosirius red stained CDMs were imaged on a Leica DM 4000 microscope at 20x magnification. Picrosirius red staining was quantified from brightfield images on ImageJ. Five ROIs per matrix were analysed with three matrices per repeat for three independent repeats. Collagen birefringence was quantified using the ImageJ Picrosirius Red Birefringence Analyser macro (https://github.com/TCox-Lab) [66].

##### Second Harmonic Generation Imaging

Second harmonic generation (SHG) imaging was performed on formalin fixed CDMs and organotypic matrices. Images were acquired on an inverted Leica DMI 6000 SP8 multiphoton microscope (Leica Biosystems) with a Ti-Sapphire femtosecond laser (Coherent Scientific) excitation source tuned to a wavelength of 880nm. SHG signal was recorded with RLD HyD filters 440/20nm. An image area of 512×512 pixels was acquired, using a bidirectional scan with a speed of 400Hz and line averaging of 4. Peak SHG intensity was quantified using an in-house quantification tool previously used [35, 65, 66]. Overall, three ROIs per matrix/well were imaged with three matrices per condition for three independent repeats.

##### Western Blotting

Cells were seeded at a density of 7.5×10^5^ cells for PyMT cancer cells and CAFs, and 9×10^5^ for MDA-MB-231 cells and human CAFs in 10cm TC treated plastic dishes. Following 24 hours of growth cells were treated with 5µM GV101 or DMSO control for one hour for PyMT cancer and MDA-MB-231 cells, 6 hours for PyMT CAF and 24 hours for human CAF cells before lysate extraction. Dishes were placed on ice and washed twice with ice-cold PBS. Following removal of final PBS wash ice-cold RIPA buffer (50nM HEPES, 1% Triton X-100, 0.5% Sodium deoxycholate, 0.1% SDS, 0.5mM EDTA, 50mM NaF, 10mM Sodium orthovanadate, 1x protease inhibitor cocktail [Complete™ mini, EDTA free, Roche]) was added to cells and incubated on ice for 10 minutes. Lysates were removed from plates with a cell scraper and centrifuged at 13,000 RPM at 4°C for 10 minutes. Supernatant was transferred into fresh Eppendorf tubes on ice and either frozen and stored at −20°C or protein quantification performed. Protein concentration was performed via a Bradford assay (Bio-Red Protein Assay Dye Reagent Concentrate). Lysates were prepared at a final concentration of 2µg/µL in sample buffer (NuPage™ LDS sample buffer x4) containing 50mM 1,4-Dithiothreitol (DTT, Invitrogen) which were subsequently denatured by heating at 70°C for 10 minutes. Samples were stored at −20°C and thawed at room temperature for western blotting.

Protein lysates (20µg) were separated via gel electrophoresis on a 4-12% Bis-Tris gel (NuPage™, Thermo Fisher Scientific). Following separation proteins were transferred onto a PVDF membrane (Immobilon-P, Millipore) in ice-cold transfer buffer (1.05% glycine, 0.225% Tris, 10% methanol in H_2_O) for one hour. Membranes were then blocked in 5% bovine serum albumin (BSA) for one hour at room temperature on a shaking platform. Membranes were then incubated in primary antibody in Tris-buffered saline (0.1% Tween):1% BSA overnight at 4°C on a shaking platform. Following incubation primary antibody was removed and membranes were washed 5x for 5 minutes each in TBST to remove any unbound antibody. Washed membranes were then incubated in horseradish peroxidase (HRP)-linked secondary antibody for 2 hours at room temperature on a shaking platform before removal and again washed 5x for 5 minutes each in TBST to remove any unbound antibody. DAB signal was detected using ECL Plus reagent for GAPDH or Ultra-Enhanced Chemiluminescence for all other antibodies (Western Lightening ECL, PerkinElmer) on a Fusion FX (Viber) machine. Densitometry analysis was performed on ImageJ and normalised to GAPDH loading control.

##### Analysis of TCGA and METABRIC datasets

TCGA mRNA expression (RNA seq V2 RSEM) and patient clinical data was downloaded for 1040 patients with breast ductal, lobular, mixed or breast carcinoma. TNBC patients were defined as ER, PR and HER2 negative on IHC. For survival analysis, patients diagnosed as stage I-III were included (999 patients). Patients were separated based on ROCK2 expression levels split in 50% high and 50% low ROCK2 expression. METABRIC ROCK2 mRNA expression (Illumina HT-12 v3 microarray) and patient clinical data was downloaded for 1936 patients with invasive breast ductal, lobular or mixed carcinoma. For survival analysis only patients who were either alive at last follow-up or died of disease were included (n=1449 patients). Patients were stratified into tertiles based on ROCK2 expression.

##### Tumour microarray analysis

Two tumour microarray (TMA) cohorts, Garvan/Royal Prince Alfred Hospital (RPAH) Breast Progression Series and RPAH/Concord Repatriation General Hospital (CRGH) TNBC cohort, were obtained for ROCK2 immuno-histochemistry (IHC) staining and analysis. Both TMA cohorts were graded and reviewed by a trained breast pathologist at time of construction. Analysis was performed using Qupath image analysis software. A total of 130 patients were used in the analysis of the Garvan/RPAH Breast Progression series and 123 patients in the RPAH/CRGH TNBC cohort. For survival analysis patients were stratified based on high (50%) or low (50%) ROCK2 stain coverage and/or intensity.

## Supplementary Figure legends

Supplementary Figure S1: GV101 reduces ROCK2 signalling and collagen remodelling by PyMT and human CAFs, related to Figure 1

**(A)** Representative western blot images showing ROCK2 protein expression in PyMT and Human TNBC CAFs. **(B)** Representative western blot images showing total and phosphorylated Cofilin (pCofilin) of PyMT CAFs following treatment with DMSO or GV101 (i) and quantification of pCofilin levels relative to total cofilin (ii). (**C)** Representative western blot images showing total and pCofilin of human TNBC CAFs following treatment with DMSO or GV101 (i) and quantification of pCofilin levels relative to total cofilin (ii). **(D-G)** Representative images of individual birefringence signal channels for picrosirius red stained PyMT CAF matrices **(D)** and quantification of birefringence signal showing contribution of green **(E)**, yellow **(F)** and red-orange **(G)** birefringence signal. **(H-K)** Representative images of individual birefringence signal channels for picrosirius red stained Human TNBC CAF matrices **(H)** and quantification of birefringence signal showing contribution of green **(I)**, yellow **(J)** and red-orange **(K)** birefringence signal to the total birefringence signal. Data represents mean ± SEM of three individual repeats. p-values were determined using a One-way ANOVA with multiple comparisons. ns = not significant (p≥0.05), * = p<0.05. Scale bar = 100µm.

Supplementary Figure S2: GV101 reduces ROCK2 signalling and cell viability in PyMT and MDA-MB-231 TNBC cell lines, related to Figure 3

**(A)** Representative western blot images showing ROCK2 protein expression in PyMT 20065 and MDA-MB-231 cancer cells. **(B)** Representative western blot images showing total and phosphorylated Cofilin (pCofilin) of PyMT cancer cells following treatment with DMSO or GV101 (i) and quantification of pCofilin levels relative to total cofilin (ii). **(C)** Representative images of western blot showing total and pCofilin of MDA-MB-231 cells following treatment with DMSO or GV101 (i) and quantification of pCofilin levels relative to total cofilin (ii). **(D)** Quantification of cell viability of PyMT cancer cells following treatment with DMSO, GV101 or Fasudil. **(E)** Quantification of cell viability of MDA-MB-231 cells following treatment with DMSO, GV101 or Fasudil. Data represents mean ± SEM of three individual repeats. p-values were determined using One-sample t test. ns = not significant (p≥0.05), * = p<0.05, ** = p<0.01.

Supplementary Figure S3: Analysis workflow for cell apoptosis quantification following exposure to fluid flow-induced shear stress, related to Figure 3 and STAR Methods

**(A)** Cellpose was used to segment cells in phase images and generate a cell coverage mask. **(B)** Matched AnnexinV masks were generated and superimposed on the corresponding cell coverage masks to determine AnnexinV signal per individual cell.

## Supplementary Movie legends

Supplementary Movie 1: GV101 reduces remodelling of pre-existing collagen by PyMT CAFs, related to Figure 1

Z-stacks of SHG signal through PyMT CAF-contracted collagen matrices (Day 12 of contraction) following treatment with DMSO, GV101 or Fasudil. Scale bar = 150µm.

Supplementary Movie S2: GV101 reduces remodelling of pre-existing collagen by human TNBC CAFs, related to Figure 1

Z-stacks of SHG signal through human CAF-contracted collagen matrices (Day 12 of contraction) following treatment with DMSO, GV101 or Fasudil. Scale bar = 150µm.

Supplementary Movie S3: GV101 reduces remodelling of *de novo* synthesised collagen by PyMT CAFs, related to Figure 2

Z-stacks of SHG signal through PyMT CAF-generated CDMs (Day 7 of CDM generation) following treatment with DMSO, GV101 or Fasudil. Scale bar = 150µm.

Supplementary Movie S4: GV101 reduces remodelling of *de novo* synthesised collagen by PyMT CAFs, related to Figure 2

Z-projections of SHG signal through PyMT CAF-generated CDMs (Day 7 of CDM generation) following treatment with DMSO, GV101 or Fasudil. Scale bar = 30µm.

Supplementary Movie S5: GV101 reduces remodelling of *de novo* synthesised collagen by human TNBC CAFs, related to Figure 2

Z-stacks of SHG signal through human CAF-generated CDMs (Day 7 of CDM generation) following treatment with DMSO, GV101 or Fasudil. Scale bar = 150µm.

Supplementary Movie S6: GV101 reduces remodelling of *de novo* synthesised collagen by human TNBC CAFs, related to Figure 2

Z-projections of SHG signal through human CAF-generated CDMs (Day 7 of CDM generation) following treatment with DMSO, GV101 or Fasudil. Scale bar = 30µm.

Supplementary Movie S7: GV101 reduces single-cell migration of MDA-MB-231 cells on CDMs, related to Figure 4

Time-lapse movies of MDA-MB-231 single-cell migration on CDMs over 12 hours in the presence of DMSO or GV101. Scale bar = 400µm.

Supplementary Movie S8: GV101-mediated inhibition of epithelial ROCK2 reduces PyMT cell attachment and spreading on CDMs without significantly affecting subsequent collective cell migration, related to Figure 5

Time-lapse movies of PyMT cell attachment and spreading on CDMs and subsequent collective cell migration over 24 hours in the presence of DMSO or GV101 (inhibition of epithelial ROCK2). Scale bar = 400µm.

Supplementary Movie S9: GV101-mediated inhibition of stromal ROCK2 during CDM generation is sufficient to reduce subsequent PyMT collective cell migration, related to Figure 5

Time-lapse movies of PyMT collective cell migration over 24 hours on CDMs that were generated in the presence of DMSO or GV101 (inhibition of stromal ROCK2). Scale bar = 400µm.

## Notes

### Competing Interest Statement

P.T. and D.H. receive reagents from Equilibre Biopharmaceuticals and Graviton Biosciences (GV101 provided free of charge). P.T. receives reagents from Kadmon Inc., InxMed (consultant) Redx Pharma, and Amplia Therapeutics. Under a licensing agreement between Amplia Therapeutics and Garvan Institute of Medical Research, K.J.M., P.T. (consultant) and D.H. are entitled to milestone payments.

### Summary of Updates

Main Manuscript and Figures updated.

