## Supplementary figures and images for "ROCK2 inhibition has a dual role in reducing ECM remodelling and cell growth, while impairing migration and invasion"

### SupplementaryFigure1

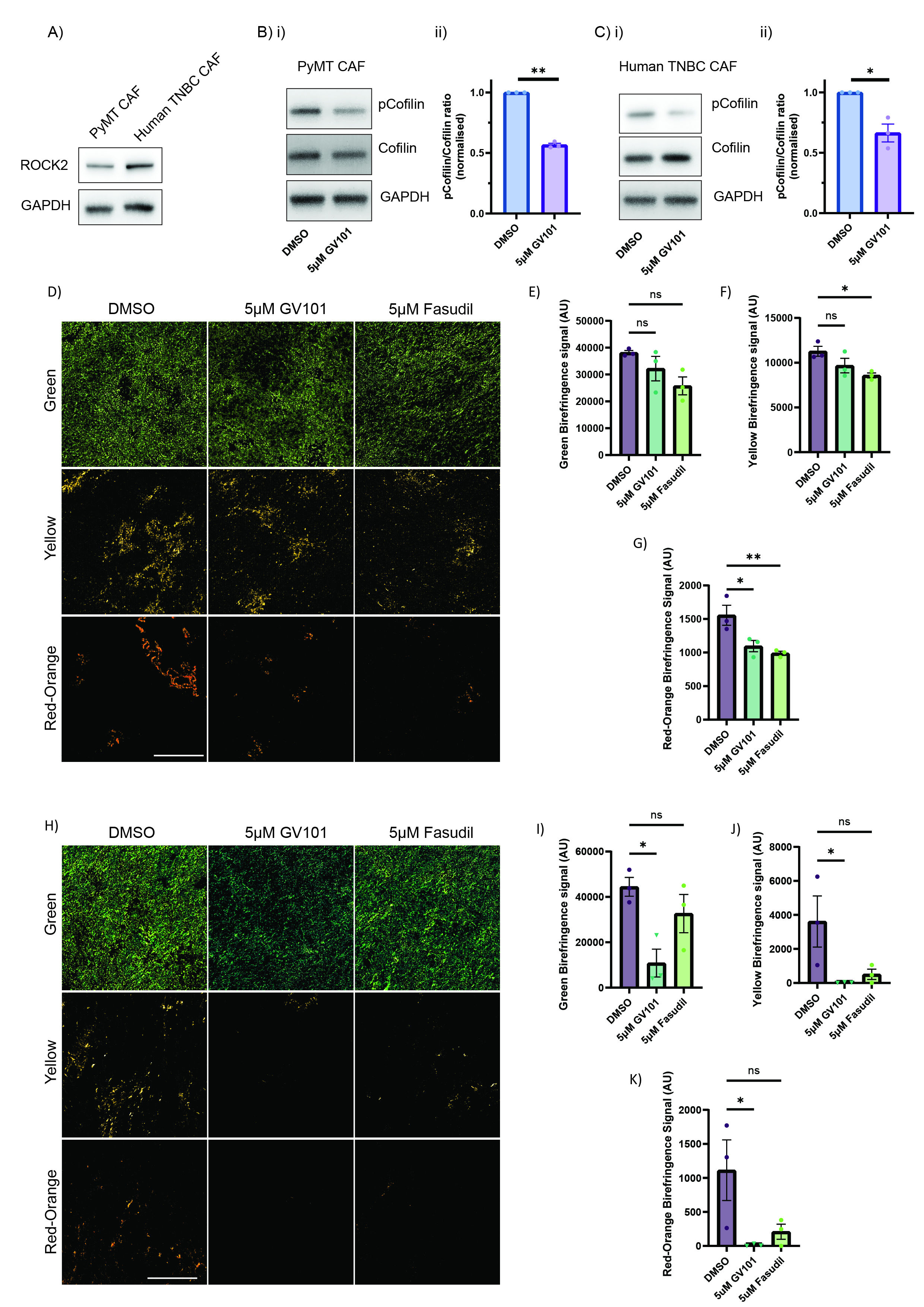

### SupplementaryFigure2

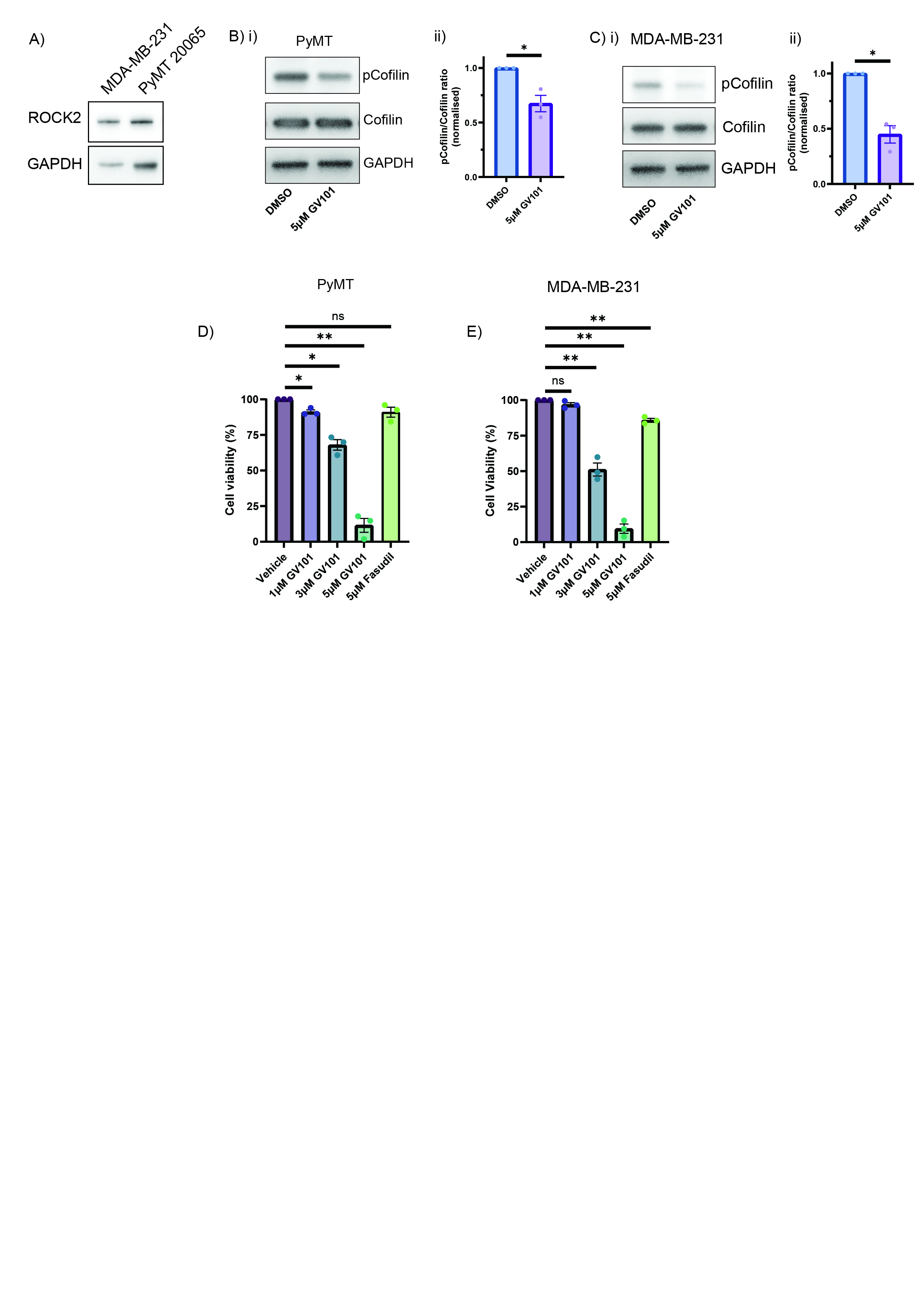

### SupplementaryFigure3

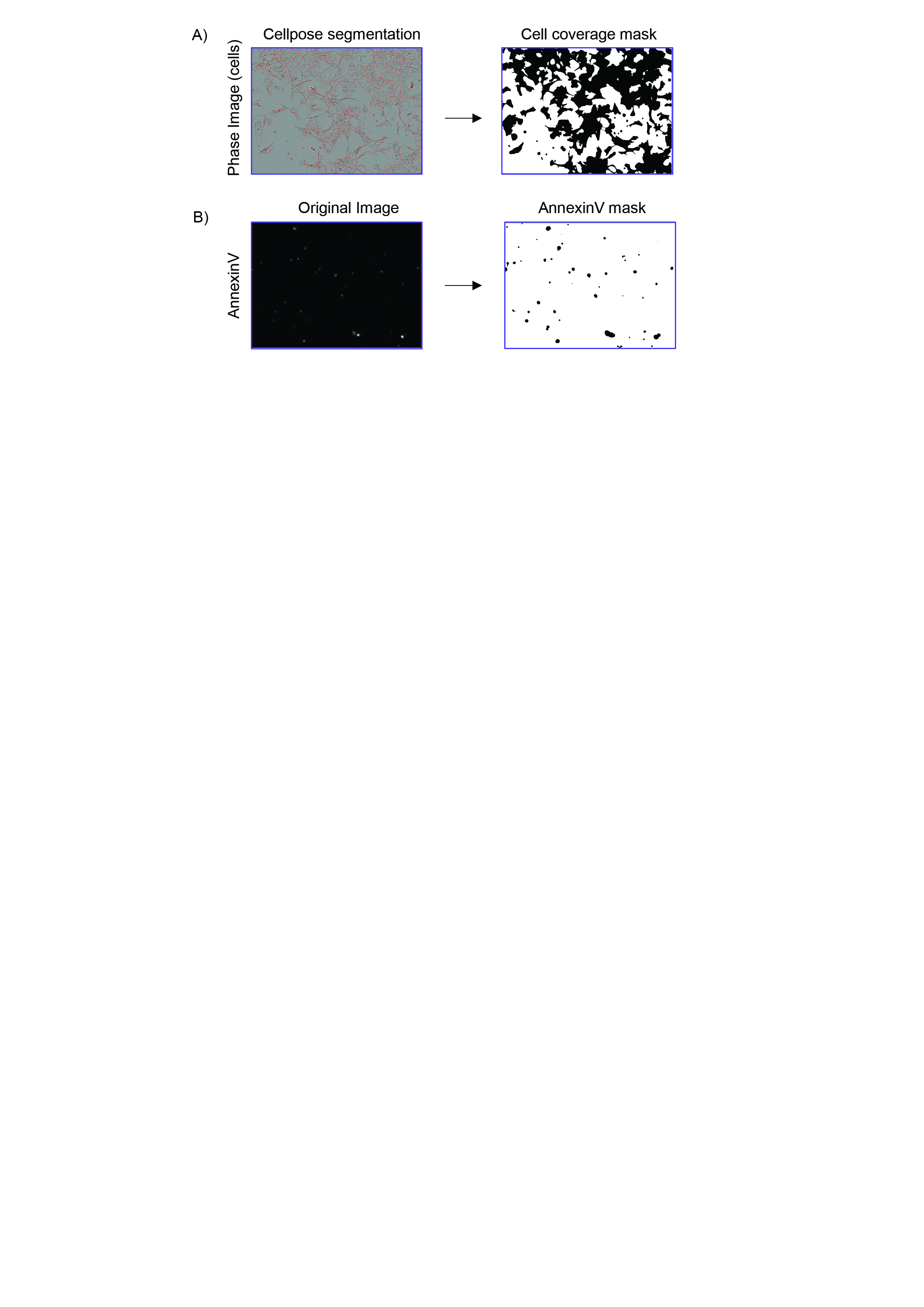
